# M1 Large-scale Network Dynamics Support Human Motor Resonance and Its Plastic Reshaping

**DOI:** 10.1101/2024.06.10.598208

**Authors:** Giacomo Guidali, Eleonora Arrigoni, Nadia Bolognini, Alberto Pisoni

## Abstract

Motor resonance – the activation of the observer’s motor system when viewing others’ actions – grounds the intertwined nature of action perception and execution, with profound implications for social cognition and action understanding. Despite extensive research, the neural underpinnings supporting motor resonance emergence and rewriting remain unexplored.

In this study, we investigated the role of sensorimotor associative learning in motor resonance neural mechanisms. To this aim, we applied cross-systems paired associative stimulation (PAS) to induce novel visuomotor associations in the human brain. This protocol, which repeatedly pairs transcranial magnetic stimulation (TMS) pulses over the primary motor cortex (M1) with visual stimuli of actions, drives the emergence of an atypical, PAS-conditioned motor resonance response. Using TMS and electroencephalography (EEG) co-registration during action observation, we tracked the M1 functional connectivity profile during this process to map the inter-areal connectivity profiles associated with typical and PAS-induced motor resonance phenomena.

Besides confirming, at the corticospinal level, the emergence of newly acquired motor resonance responses at the cost of typical ones after PAS administration, our results reveal dissociable aspects of motor resonance in M1 interregional communication. On the one side, typical motor resonance effects acquired through the lifespan are associated with prominent M1 alpha-band and reduced beta-band connectivity, which might facilitate the corticospinal output while integrating visuomotor information. Conversely, the atypical PAS-induced motor resonance is linked to M1 beta-band cortical connectivity modulations, only partially overlapping with interregional communication patterns related to the typical mirroring responses. This evidence suggests that beta-phase synchronization may be the critical mechanism supporting the formation of motor resonance by coordinating the activity of motor regions during action observation, which also involves alpha-band top-down control of frontal areas.

These findings provide new insights into the neural dynamics underlying (typical and newly acquired) motor resonance, highlighting the role of large-scale interregional communication in sensorimotor associative learning within the action observation network.

## 1. INTRODUCTION

Motor resonance is a visuomotor matching mechanism endowed in the human brain, whereby observing an action covertly activates the corresponding motor programs in the observer’s central nervous system (Rizzolatti & Craighero, 2004). This phenomenon can be measured by recording motor-evoked potentials (MEPs) induced by transcranial magnetic stimulation (TMS) while observing biological movements (e.g., grasping a mug), where MEPs have higher peak-to-peak amplitudes than during the observation of static stimuli (e.g., a static hand; Fadiga, Fogassi, Pavesi, & Rizzolatti, 1995). Considering simple movements, motor resonance is typically hemispheric-specific: the observation of unilateral hand movements recruits the contralateral motor system only, as in the case of their execution (Aziz-Zadeh, Maeda, Zaidel, Mazziotta, & Iacoboni, 2002). The increased reactivity of the primary motor cortex (M1) during action observation has been interpreted as evidence of the recruitment of mirror neurons within the so-called *action observation network* (AON), and it is considered one of the most reliable indexes for non-invasively investigating motor resonance in the human brain (for a review, see: Naish, Houston-Price, Bremner, & Holmes, 2014). Notably, embodied theories of cognition hold that motor resonance is necessary for understanding the intention of others and for the inference of their action goals (Gallese & Goldman, 1998; Kaplan & Iacoboni, 2006; Keysers & Gazzola, 2006), and experimental evidence suggests a pivotal role of this phenomenon for social cognition, action understanding, and emphatic abilities (e.g., Avenanti, Sirigu, & Aglioti, 2010; Bekkali et al., 2021; Bucchioni, Cavallo, Ippolito, Marton, & Castiello, 2013; Guidali, Picardi, Franca, Caronni, & Bolognini, 2023; Hogeveen & Obhi, 2012; Lo Gerfo et al., 2018; Mehta, Ashok, Thirthalli, & Keshavan, 2019; Orban, Lanzilotto, & Bonini, 2021; Pisoni et al., 2014).

However, a still debated question is how visuomotor associations grounding motor resonance are formed, giving rise to AON activation (Heyes & Catmur, 2022). Phylogenetic accounts suggest that mirror mechanisms have evolved genetically as an adaptation of visuomotor neurons for understanding actions (e.g., Bonini & Ferrari, 2011; Rizzolatti & Arbib, 1998), whereas ontogenetic ones suggest that mirror neurons acquire their cross-modal properties through repeated sensorimotor experiences (e.g., Catmur, Press, & Heyes, 2016; Cook, Bird, Catmur, Press, & Heyes, 2014). Considering the latter, a plausible mechanism contributing to the emergence of motor resonance during development involves the repeated direct mapping between the visual representations of others’ actions (i.e., action observation) and one’s own motor representations (i.e., action execution), which, in turn, could shape humans’ mirroring responses to observed actions (Catmur, 2013; Heyes, 2010; Keysers & Gazzola, 2014). Accordingly, several studies demonstrated how AON activation, and hence motor resonance responses, can be modulated by exploiting visuomotor paradigms repeatedly coupling the observation and the execution of a target movement, as well by coupling M1 exogenous stimulation with action observation, suggesting a crucial role of sensorimotor associative learning not only in the development of the AON but also in its plastic rewriting during adulthood (Bardi, Bundt, Notebaert, & Brass, 2015; Brunsdon, Bradford, Smith, & Ferguson, 2020; Catmur, Walsh, & Heyes, 2007; de Klerk, Johnson, Heyes, & Southgate, 2015; de Klerk, Lamy-Yang, & Southgate, 2019; Fitzgibbon et al., 2016; Guidali, Carneiro, & Bolognini, 2020; Guidali, Picardi, Gramegna, & Bolognini, 2023).

Despite the compelling literature on mirroring mechanisms during action observation (for a review, see: Kemmerer, 2021), little is known about the large-scale network dynamics supporting motor resonance and its plastic modulation. The human AON has been described as relying upon different cortical areas densely interconnected, encompassing, beyond M1, frontal (e.g., inferior frontal gyrus, ventral premotor cortex, supplementary motor cortex), parietal (e.g., superior and inferior parietal lobule), and temporal regions (e.g., middle temporal gyrus) (Caspers, Zilles, Laird, & Eickhoff, 2010; Molenberghs, Cunnington, & Mattingley, 2012). Given the distinct functional roles of these areas within the AON (e.g., Caspers et al., 2010), the interregional dynamics governing visuomotor associations underlying motor resonance and its occurrence are still unknown. For instance, a still open question is whether typical (i.e., acquired through the lifespan) and newly acquired (i.e., experimentally induced) motor resonance responses are characterized by the same cortico-cortical patterns while presenting the same peripheral output (i.e., MEP facilitation during action observation), or else if these mechanisms change through experience and thus result different in time. Hence, investigating these phenomena at a whole-brain level is essential to better characterize motor resonance features, their emergence, and, in a broader perspective, the global dynamics of brain networks with mirror properties.

Recently, our research group introduced a *cross-systems* paired associative stimulation (PAS) protocol that proved to be effective in creating novel motor resonance responses to conditioned visual stimuli of actions, likely through the induction of associative plasticity within the AON: the mirror PAS (m-PAS) (Guidali et al., 2020). PAS are non-invasive brain stimulation protocols that induce Hebbian-like associative plasticity through the repeated pairing of cortical and peripheral stimulation; the former is delivered with TMS, and the latter occurs through different sensory modalities (Kraus et al., 2018; Ranieri et al., 2019; Schecklmann et al., 2011; Sowman, Dueholm, Rasmussen, & Mrachacz-Kersting, 2014; Stefan, Kunesch, Cohen, Benecke, & Classen, 2000; Suppa et al., 2013; Suppa, Li Voti, Rocchi, Papazachariadis, & Berardelli, 2015; Wolters et al., 2005; Zazio, Guidali, Maddaluno, Miniussi, & Bolognini, 2019) (for a review, see: Guidali, Roncoroni, & Bolognini, 2021). Specifically, in the m-PAS, TMS pulses are repeatedly delivered over the right M1 coupled with the observation of visual stimuli depicting a finger movement made with the ipsilateral (with respect to TMS site of stimulation) right hand, with a specific temporal delay based on the chronometry of the brain mechanisms responsible for motor control (i.e., 25 ms; Guidali et al., 2020). At baseline, this visual stimulus does not induce motor resonance in the ipsilateral hemisphere. Nevertheless, after m-PAS administration, the successful induction of atypical motor resonance emerges at a neurophysiological level, as indexed by corticospinal facilitation in the ipsilateral hemisphere when the PAS-conditioned movement is observed (Guidali et al., 2020). This neurophysiological effect has a behavioral counterpart: the automatic imitation of the PAS-conditioned action is facilitated accordingly (Guidali, Picardi, Gramegna, et al., 2023). So, the m-PAS offers a promising benchmark to investigate the effects of visuomotor associative plasticity within the AON, shedding light on the role of large-scale interregional communication mediating mirror activations during action observation and their development.

Starting from this premise, the present study explores motor resonance and its plastic side by taking advantage of TMS and electroencephalography (EEG) co-registration. TMS-EEG enables the real-time exploration of cortical reactivity at a whole-brain level by examining TMS-evoked potentials (TEPs). TEPs are considered a reliable indicator of cortical excitability, offering insights into the activation state of the stimulated region (Casarotto et al., 2010; de Tommaso et al., 2020; Miniussi & Thut, 2010). Crucially, the analysis of TEPs allows us to observe how the activity triggered by TMS propagates over time and space, reaching distant regions connected through functional pathways (Bortoletto, Veniero, Thut, & Miniussi, 2015; Pisoni, Mattavelli, et al., 2018; Rogasch & Fitzgerald, 2013). Examining the oscillatory dynamics of TEPs can then offer insights into inter-regional functional connectivity, potentially shedding light on communication between different brain areas (Bianco, Arrigoni, Di Russo, Romero Lauro, & Pisoni, 2023; Pisoni, Romero Lauro, Vergallito, Maddaluno, & Bolognini, 2018; Thut & Miniussi, 2009). Finally, TMS-EEG has been extensively used to characterize the connectivity dynamics of the motor system in healthy and pathological conditions (Hernandez-Pavon et al., 2023).

By taking advantage of TMS-EEG co-registration, our aim is twofold: (*a*) to better characterize the cortico-cortical signatures of typical and newly acquired motor resonance responses and, by doing this, (*b*) to shed light on the neurophysiological fingerprints of sensorimotor learning and associative plasticity within the AON, using the m-PAS protocol as an operative model for motor resonance rewriting. Studying large-scale dynamics could better ground the neurophysiological bases of (typical and induced) motor resonance, as well as of sensorimotor associative learning within mirror networks. By analyzing inter-areal communication, we can describe the functional networks related to motor facilitation and how these differ with newly acquired visuomotor association transiently induced by the m-PAS protocol. We hypothesize that these plastic changes encompass a distributed network of cortical regions, reflecting important AON nodes connected to the stimulated right hemisphere’s M1 and only partially overlapped with the canonical ones.

Healthy participants underwent an action-observation paradigm before and after administering the m-PAS protocol (Guidali et al., 2020). During the task, static or moving index finger of the left (i.e., contralateral to the stimulated M1 – to evaluate the typical motor resonance response) and right hand (i.e., ipsilateral to the stimulated M1 – to assess the atypical response, potentially induced during the m-PAS) were visually presented. We recorded TEPs from the right M1 during this task to assess cortical connectivity patterns. We focused on the source-level functional connectivity profile of M1 in typical and newly acquired motor resonance conditions across alpha (8-12 Hz) and beta (13-30 Hz) frequency bands, which are known to be crucial for the sensorimotor system activity during action observation (e.g., Babiloni et al., 2016; Muthukumaraswamy & Johnson, 2004; Qin et al., 2023; Simon & Mukamel, 2016). Simultaneously, we recorded MEPs from two muscles of the left hand (i.e., *first dorsal interosseus* – FDI – and *abductor digiti minimi* – ADM) to investigate motor resonance at the corticospinal level and control for the protocol’s effectiveness.

## 2. METHODS and MATERIALS

### 2.1. Participants

Twenty-five healthy volunteers (10 males, mean age ± standard deviation – SD = 24.5 ± 2.2 years; mean education ± SD = 16.6 ± 2 years) with no neurologic problems or contraindications to TMS took part in the study. We determined the sample size through an a-priori within-subjects repeated measures analysis of variance (rmANOVA) using the software G*Power 3.1 (Faul, Erdfelder, Buchner, & Lang, 2009) and selecting as the target effect size value the smallest one among those found in our previous studies using the m-PAS and exploiting corticospinal motor resonance (Guidali et al., 2020; Guidali, Picardi, Gramegna, et al., 2023). This value corresponded to an partial eta squared (*η_p_^2^*) of .16 (see Experiment 1 results in Guidali, Picardi, Gramegna, et al., 2023). The power analysis (alpha error level: *p* = .05; statistical power = .9, actual power = .93) showed a recommended sample size of at least 24 participants to achieve enough statistical power. According to the Edinburgh Handedness Inventory (Oldfield, 1971), all participants were right-handed (mean score ± SD = 78.4 ± 12.1%), and none had contraindications to TMS (Rossi et al., 2021). The experiment was performed following the ethical standards of the Declaration of Helsinki after the approval of the Ethical Committee of the University of Milano-Bicocca. Before taking part in the study, participants gave their written informed consent. Tasks’ scripts, raw data, and analysis of this study will be publicly available at Open Science Framework (OSF) once the work will be accepted on a peer-review journal.

### 2.2. Experimental procedure

The study comprised one TMS-EEG session, lasting about 2 hours and a half, in which participants underwent the standard m-PAS protocol, whose effectiveness was already replicated in two previous studies (Guidali et al., 2020; Guidali, Picardi, Gramegna, et al., 2023). The session started with the preparation of the EEG cap and EMG electrodes. Then, the experimenters determined the participant’s motor hotspot of the left hand’s FDI muscle and assessed the individual resting motor threshold (rMT). Subsequently, the blocks of the action observation task were administered. Following this task, participants underwent the m-PAS, and immediately after its end, motor resonance was re-assessed with the action observation task (**Figure 1**).

**Figure 1.**
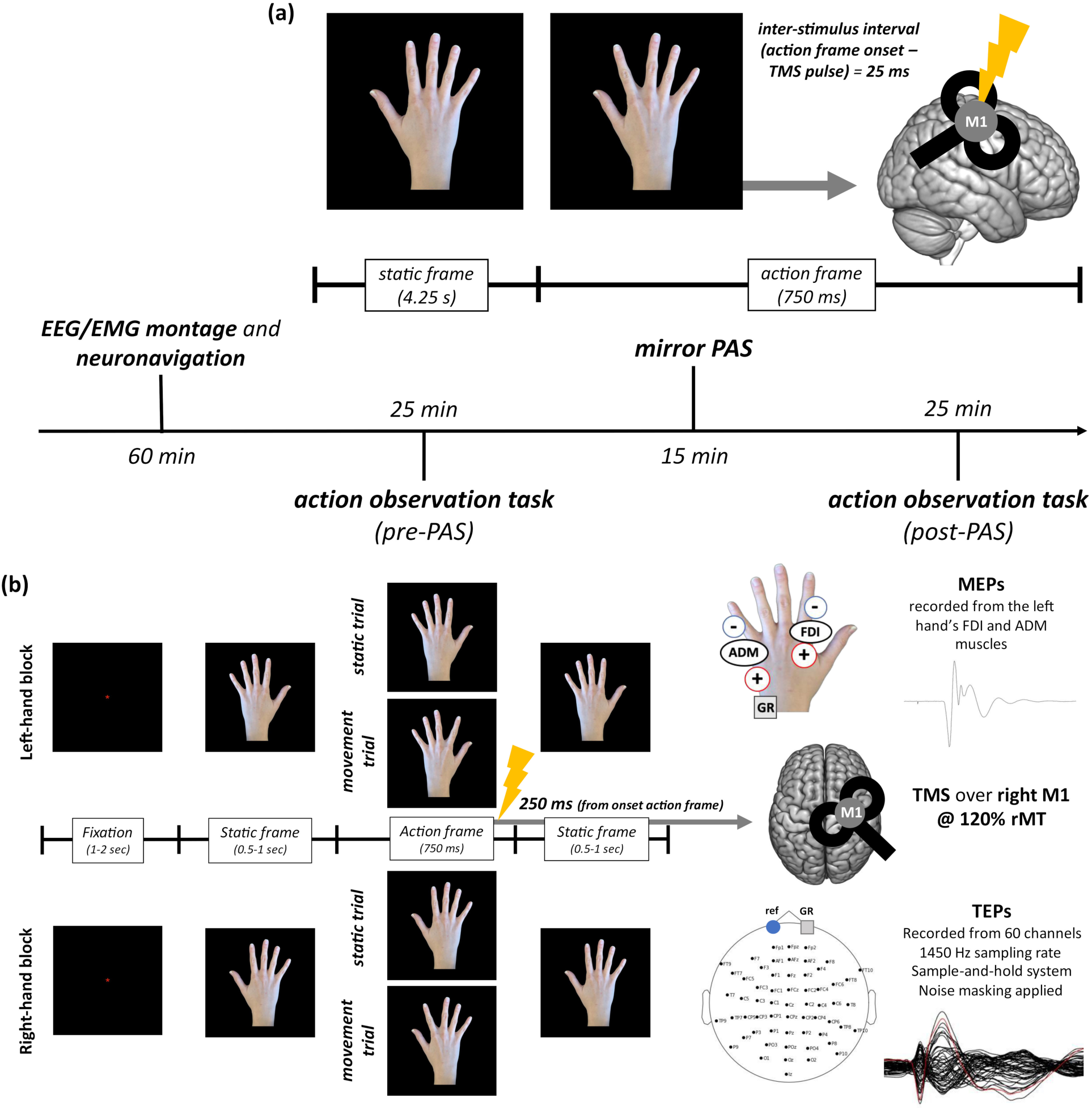
Experimental procedure of the study. After the montage of the EEG cap and left hand’s EMG electrodes, participants underwent the blocks of the action observation task (**b**) before and after administering the m-PAS (**a**). m-PAS parameters were taken from the effective ones found in our previous studies (Guidali et al., 2020; 2023). The action observation task blocks comprised trials depicting a static/moving left-hand or right-hand index finger (for a similar procedure, see: Guidali et al., 2020). Throughout the experiment, TMS was always delivered over the right M1. MEPs were recorded from two muscles of the left hand: i.e., FDI (target muscle, involved in the observed movement) and ADM (control muscle, not involved in the observed movement).

### 2.3. m-PAS protocol

The m-PAS protocol is a visuomotor, *cross-systems* PAS protocol in which the peripheral stimulation paired with TMS over the right M1 is a visual stimulus depicting the abduction/adduction movement of the right-hand index finger (Guidali et al., 2020). During the protocol, participants sat comfortably on a chair in front of a PC monitor at a distance of 100 cm with their hands positioned out of view on the chair’s armrests. Each trial of the m-PAS began with a frame (i.e., ‘static frame’) depicting the dorsal view of the right hand in an egocentric perspective for a fixed duration of 4250 ms. Immediately after its end, a second frame (‘action frame’) appeared, lasting 750 ms and showing the abduction movement of the index finger of the same right hand (**Figure 1a**). The rapid succession of the ‘static’ and ‘action’ frames gave the illusion of apparent motion. After 25 ms from the ‘action frame’ onset, a TMS pulse was delivered over the right M1 at 120% of the participant’s rMT. The timing of the stimuli was presented under computer control (E-Prime 2.0, Psychology Software Tool, Inc.), and the frames’ timing was checked using a photodiode before data collection. One hundred and eighty trials were presented at a frequency of .2 Hz for 15 min. To control that participants were looking with attention to the visual stimuli, a critical condition for the success of a PAS protocol (Stefan, Wycislo, & Classen, 2004), in 15 trials out of 180, a red circle appeared on the fingernail of the moving index finger. Participants were instructed to press as fast and accurately as possible with their right hand on the left key of the PC mouse as soon as the circle appeared. On average, participants’ accuracy in this task was 98.1% (SD = ± 2.94%).

### 2.4. Action observation task

Corticospinal and cortical excitability during the observation of movements were measured before and after m-PAS administration, recording MEPs and TEPs induced by the stimulation of the right M1 during a standard action observation task. Here, participants observed visual stimuli showing static or moving hand stimuli (same task as in: Guidali et al., 2020; Guidali, Picardi, Gramegna, et al., 2023). MEPs were recorded from two muscles of the left hand: i.e., FDI (target muscle, involved in the observed movement) ADM (control muscle, not involved in the observed movement). MEPs and TEPs were simultaneously collected. Every trial began with a fixation point (a red asterisk) presented on the screen’s black background (duration jittered between 1000 and 2000 ms). As soon as the fixation disappeared, the dorsal view of a right/left hand was presented in an egocentric perspective and centrally on the screen (the same posture as the ones depicted during the m-PAS), for a variable duration from 800 to 1000 ms. Then, a third frame appeared (fixed duration of 750 ms). In ‘movement trials’, the frame showed the abduction movement of the index finger (the same movement shown during the m-PAS). In ‘static trials’, the previous frame still showed, and thus the hand remained static. In both kinds of trials, 250 ms after the onset of this third frame, a TMS pulse was delivered over the right M1. TMS intensity was set at 120% of the participant’s rMT. Finally, the frame depicting the static hand reappeared for 750 ms, and the trial ended (**Figure 1b**). Four blocks of trials, two showing a left hand and the others showing a right hand, were presented. In each block (lasting about 5 min), 80 trials were presented in a randomized order: half (40) of the trials showed the static hand, and the other half (40) had the moving index finger. The order of the blocks was counter-balanced among participants, and blocks depicting the same hand were never presented consecutively. Participants made a brief pause (approximately 1 min) between one block and the other. To ensure that participants kept attention to the visual stimuli, in each block of the action-observation task, 4 trials out of 80 presented a small (diameter: 15 pixel) colored circle that appeared on the fingernail of the index finger or the middle finger (in a randomized order) during the third frame of the trial. Participants had to press the mouse with their right hand every time the dot appeared. On average, participants’ accuracy in this attentive task was 95.2% (SD = ± 2.7%). TEPs and MEPs recorded during these attentional trials were not analyzed. Trials randomization and timing of the stimuli were presented under computer control (E-Prime 2.0, Psychology Software Tool, Inc.).

### 2.5. TMS

TMS pulses were delivered with a biphasic figure-of-eight coil (diameter = 70 mm) connected to a Nexstim Eximia stimulator (Nexstim, Helsinki, Finland). At the beginning of the session, the motor hotspot of the left-hand FDI muscle was found by moving the coil around the motor hand area (i.e., hand knob) of the right hemisphere by using a slightly supra-threshold stimulus and recording MEPs. The stimulation target site was localized on participants normalized high-resolution magnetic resonance images – MR. The individual rMT was calculated using the parameter estimation by sequential testing (PEST) procedure, a maximum-likelihood threshold-hunting procedure optimized for rMT detection (Awiszus, 2003; Dissanayaka, Zoghi, Farrell, Egan, & Jaberzadeh, 2018). On average, participants presented an rMT of 42.8% (SD = ± 8.2%). After the determination of rMT, we ensure that an early cortical response (30 ms) of at least 5 μV was recorded at the target experimental intensity (i.e., 120% rMT) during the stimulation of the right M1, as assessed online in a short recording session before the experimental blocks. The stable TMS coil placement and position were constantly monitored during the experimental sessions using the stimulator’s integrated Navigated Brain Stimulation (NBS) system (Nexstim™, Helsinki, Finland) based on infrared-based frameless stereotaxy. The software allows accurate monitoring of the coil position and orientation as well as online estimation of the distribution and intensity (V/m) of the intracranial electric field induced by the TMS (mean electric field induced at 120% rMT ± SD = 87.4 ± 19.1 V/m). The coil was positioned tangentially to the scalp and tilted 45° to the midline (positioned perpendicular to the stimulated cortical gyrus), inducing currents in the brain with an anterior-to-posterior (first phase) / posterior-to-anterior (second phase) direction. TMS was always administered with these parameters throughout the session.

### 2.6. EMG recordings

MEPs were recorded from right-hand FDI and ADM muscles using Signal software (version 3.13) connected to a Digitmer D360 amplifier and a CED micro1401 A/D converter (Cambridge Electronic Devices, Cambridge, UK, www.ced.co.uk). Active electrodes (15 × 20 mm Ag-AgCl pre-gelled surface electrodes, Friendship Medical, Xi’an, China) were placed over the muscle bellies; reference electrodes over the metacarpophalangeal joint of the index finger for FDI and the little finger for ADM. The ground electrode was placed over the left head of the ulna. Before data acquisition, a visual check by the experimenters guaranteed that background noise did not exceed 20 μV. EMG signals were sampled (5000 Hz), amplified, band-pass filtered (10–1000 Hz) with a 50 Hz notch filter, and stored for offline analysis. Data was collected from 100 ms before to 200 ms after the TMS pulse (time window width: 300 ms).

### 2.7. EMG preprocessing

MEPs were analyzed offline using Signal software (version 3.13), exploiting the standard preprocessing pipeline used in our laboratory (Guidali, Picardi, Gramegna, et al., 2023). At first, trials with muscular artifacts or background noise deviating from 200 µV in the 100 ms before the TMS pulse were automatically excluded from the analysis. Then, MEP peak-to-peak amplitude was calculated in each trial in the time window between 5 ms and 60 ms from the TMS pulse. We excluded from the following analyses trials in which MEP amplitude was smaller than 50 µV. Then, for each session’s time point (i.e., pre-PAS, post-PAS), trials where MEPs amplitude were ± 2 SD from the mean of each condition (i.e., static trials, movement trials) were considered outliers and excluded from analysis. On average, for each participant, and considering all our criteria, the 3.73% (SD = ± 5.2%) of MEPs recorded were discarded.

### 2.8. EEG recordings

EEG data was continuously recorded using a 60-channel EEG cap (EasyCap, BrainProducts GmbH, Munich, Germany) with a sample-and-hold TMS-compatible system (Nexstim™, Helsinki, Finland). Two electrodes were placed over the forehead as ground and reference. Additionally, two electro-oculographic (EOG) channels were placed over the right eyebrow and the left cheekbone to detect and monitor ocular artifacts associated with eye movements and blinking. Noise-masking was implemented by playing a custom audio track generated by randomizing TMS discharge noise through earplugs to prevent the emergence of auditory evoked potentials (Bianco et al., 2023). Individual adjustments were made to ensure the noise-masking covered TMS clicks effectively, and the volume was kept fixed throughout the experimental session. Electrodes’ impedance was kept below 5 kΩ, and the EEG signal was acquired with a sampling rate of 1450 Hz.

### 2.9. EEG preprocessing

EEG data preprocessing was performed with MATLAB R2019a (Mathworks, Natick, MA, USA). First, trials containing excessive ocular, muscle, or magnetic artifacts were detected by visual inspection and excluded. Bad channels were marked and subsequently interpolated through the spherical interpolation function of EEGLAB (Delorme & Makeig, 2004). Data were downsampled at ½ of the sampling rate (i.e., 725 Hz). Then, a notch filter between 49 and 51 Hz and a band-pass filter between 2 Hz and 50 Hz were applied. Data were segmented into epochs spanning from –1000 ms before to 1000 ms after the onset of the TMS pulse, average re-referenced, and baseline corrected for the interval between –575 and –275 ms before the TMS pulse in order to avoid the event-related activity due to stimuli visual processing (Pisoni, Romero Lauro, et al., 2018). Finally, INFOMAX Independent Component Analysis (ICA) was used to systematically remove residual magnetic artifacts, activity related to muscle contraction, blinks, and eye movements. Cleaned epochs were used to compute TEPs for each experimental condition (means ± SD – pre-PAS = left-hand static: 74.4 ± 4.1; left-hand movement: 73.5 ± 5.8; right-hand static: 75.4 ± 3; right-hand movement: 74.4 ± 4.9; post-PAS = left-hand static: 74.6 ± 3.7; left-hand movement: 72.8 ± 6; right-hand static: 73.5 ± 4; right-hand movement: 73.4 ± 4.3; all *p*s > .05). Sensor level grand average of the TEPs recorded in the different experimental conditions are reported in the **Supplementary Figure 1**.

### 2.10. Source reconstruction

TMS-evoked responses were reconstructed at the source level. We computed the forward model from a Boundary Element Model (BEM) derived by segmenting a subject’s MRI into five standard tissues (i.e., gray and white matter, cerebrospinal fluid, skull, and scalp). Fixed, standard conductivity values were assigned for the scalp, skull, and brain compartments based on the literature (Fuchs, Kastner, Wagner, Hawes, & Ebersole, 2002; Vorwerk et al., 2014). Cortical reconstruction and volumetric segmentation of the gray matter was executed using Freesurfer (Fischl, 2012), downsampled to 8194 cortical sources, and subsequently realigned to the head model space. Individual lead-field matrices were generated by aligning the forward model with the individual digitized electrode positions. Exact low-resolution brain electromagnetic tomography (Pascual-Marqui et al., 2011) was used to compute the inverse solution. The EEG source-reconstructed time series were then collapsed onto 88 regions of the automated anatomical labeling (AAL) atlas (Tzourio-Mazoyer et al., 2002) and then averaged for each parcel.

### 2.11. Statistical analyses

#### 2.11.1. Motor resonance at the corticospinal level (MEPs)

To assess and replicate m-PAS effects at the corticospinal level (Guidali et al., 2020), a *motor resonance index* was computed as the ratio in MEP amplitude between movement and static trials as follows:

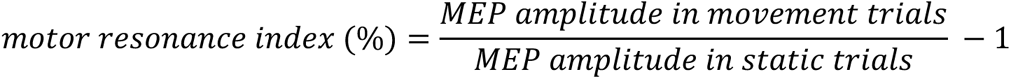

Namely, the mean MEP amplitude in movement trials was divided for MEP amplitude in static hand ones. The value ‘1’ was subtracted from the ratio to express the percentage concerning the static condition. Hence, positive values indicated corticospinal excitability facilitation by action observation (and thus the presence of motor resonance). In contrast, negative ones indicated corticospinal excitability inhibition by action observation (Guidali, Picardi, Franca, et al., 2023). We then assessed the effects of m-PAS on motor resonance with a ‘viewed Hand’ (left-hand, right-hand) X ‘Time’ (pre-PAS, post-PAS) X ‘Muscle’ (FDI, ADM) rmANOVA. MEP amplitude raw values and pre-*vs.* post-PAS planned comparisons (i.e., two-tailed paired-samples t-test) were reported in **Table 1** to assess better the contribution of static and movement trial modulation to the pattern observed on the *motor resonance index*.

**Table 1.**
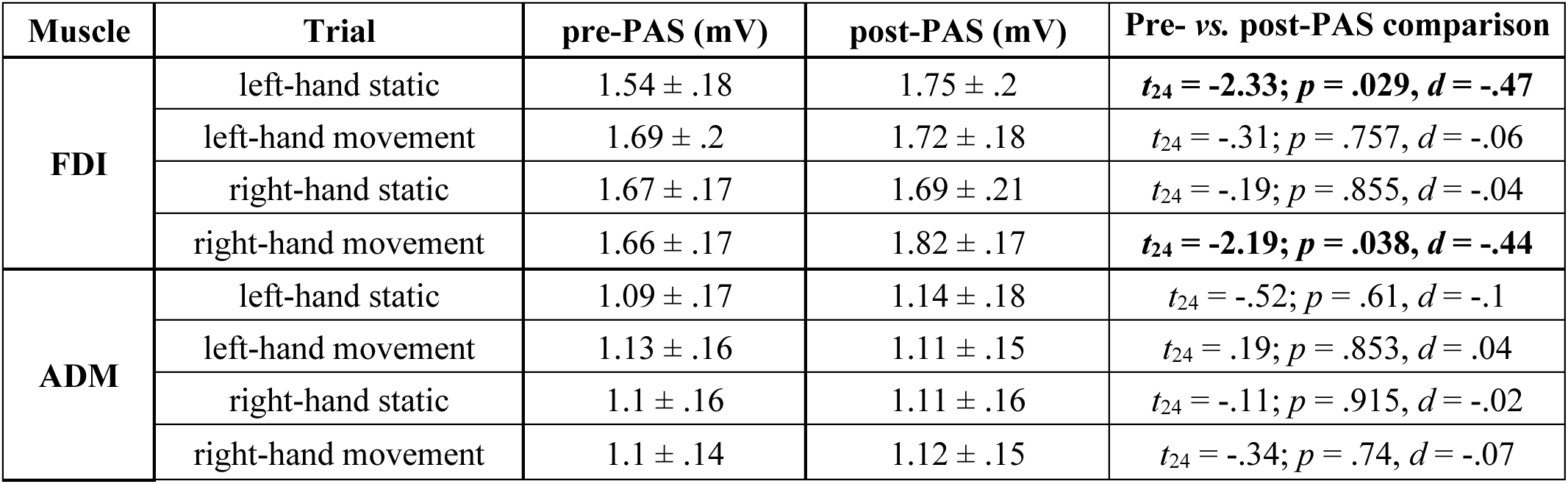
MEP amplitude raw data (mean ± standard error) from FDI and ADM muscles in the four trial typologies of the action observation task. Two-tailed paired-sample *t-tests* are reported for pre-*vs.* post-PAS comparisons. Significant ones are highlighted in bold.

#### 2.11.2. Connectivity analysis of the M1 network during action observation

In order to characterize changes in interregional dynamics induced by the m-PAS, time-frequency analysis was first conducted utilizing a multitaper method as implemented in Fieldtrip (Oostenveld, Fries, Maris, & Schoffelen, 2011). Subsequently, the debiased weighted Phase Lag Index (wPLI, Vinck et al., 2011) was calculated to assess connectivity between the right M1 and other brain parcels within the alpha (8-12 Hz) and beta (13-30 Hz) frequency bands for each task’s condition. We decided to focus our investigation on these two frequency bands, considering their prominent role in action observation and AON functioning (Babiloni et al., 2016; Muthukumaraswamy & Johnson, 2004; Qin et al., 2023; Simon & Mukamel, 2016).

To address potential spurious connectivity, we estimated wPLI in the alpha and beta band in a surrogate dataset generated by shuffling the phase of the source-reconstructed time series of each subject. This procedure was repeated for each experimental condition. A t-test was then carried out on each connectivity value representing a connection between each parcel pair (87 connections between M1 and other brain parcels), comparing real data with surrogate data to mitigate the risk of false-positive results. The correction for multiple comparisons was performed based on a 5000-permutation approach implemented in MATLAB, with a significance level set at *p* = .001 (Bianco et al., 2023). The surviving connections were eventually visualized for each condition and frequency band to highlight the resulting M1 functional connectivity patterns.

Connectivity strength was computed for each participant as the sum of all significant connections (i.e., *connectivity strength index*) and was statistically compared across the different task’s conditions with two ‘viewed Hand’ (left-hand, right-hand) X ‘Visual stimulus’ (static hand, moving hand) x ‘Time’ (pre-PAS, post-PAS) rmANOVA, one for each frequency band of interest.

Statistical analyses were performed using the software Jamovi (The Jamovi Project, 2023), R Studio (R Core Team, 2020), and Fieldtrip (Oostenveld et al., 2011). Statistical significance was set at *p* < .05, if not otherwise specified. The normality of the distributions was confirmed by checking it with the Shapiro-Wilk test and Q-Q plots assessment. All our variables were normally distributed, with the only exception being the *connectivity strength index*. Transforming this variable with the base-ten logarithm [i.e., log_10_(raw data)] made the distribution closer to normality, so this variable was analyzed using such transformation. Significant main effects and interactions were explored with multiple Bonferroni-corrected post-hoc comparisons. Partial eta-squared (*η_p_^2^*) and Cohen’s *d* were calculated in every rmANOVA and t-test, respectively, and reported as effect size values. If not otherwise specified, mean ± standard error is reported for each variable.

## 3. RESULTS

### 3.1. Effects of m-PAS on corticospinal motor resonance

Results from the rmANOVA on the *motor resonance index* (i.e., the ratio in MEP amplitude between movement and static trials) showed a significant ‘viewed Hand’ X ‘Time’ X ‘Muscle’ interaction (*F*_1,24_ = 9.8, *p* = .005, *η_p_^2^* = .29), as well as main effect of factor ‘Muscle’ (*F*_1,24_ = 6.54, *p* = .017, *η_p_^2^* = .21) and interaction ‘viewed Hand’ X ‘Time’ (*F*_1,24_ = 6.54, *p* = .017, *η_p_^2^* = .21). No other significant effect was found (all *Fs* < .46, all *ps* > .507). Corticospinal motor resonance patterns were further explored with two separate rmANOVA, one for each muscle, to understand m-PAS modulations better.

For the FDI muscle – the muscle involved in the movement observed during the m-PAS – this analysis showed significant main effects of the ‘viewed Hand’ X ‘Time’ interaction (*F*_1,24_ = 20.91, *p* < .001, *η_p_^2^* = .47). No other significant effect was found (all *F*s < .46, all *p*s > .507). Post-hoc comparisons showed that, as expected, (typical) motor resonance at baseline is present only for left-hand conditions (mean *motor resonance index* ± standard error: 10.04 ± 2.29%). Importantly, right-hand conditions did not give rise to motor resonance (–0.07 ± 1.66%; *vs.* pre-PAS left-hand *motor resonance index*: *t*_24_ = 3.61; *p* = .008, *d* = .72). A deeper look at raw MEP amplitude obtained in left-hand static and movement trials (**Table 1**) highlighted the classic enhancement of corticospinal excitability during action observation (left-hand static trials *vs.* left-hand movement trials: *t*_24_ = –4.8; *p* < .001, *d* = –.96). As expected (Guidali et al., 2020; Guidali, Picardi, Gramegna, et al., 2023), motor resonance for the PAS-conditioned right-hand movements emerged after the administration of the m-PAS (8.4% ± 1.27%; *vs.* pre-PAS right-hand *motor resonance index*: *t*_24_ = 4.54, *p* < .001, *d* = 0.91; **Figure 2a**). MEP amplitude patterns highlighted a significant enhancement of corticospinal excitability while observing movement trials, responsible for the emergence of the experimentally induced motor resonance (**Table 1**). Interestingly, the newly acquired motor resonance response was associated with a rewriting of the typical one. Indeed, after the m-PAS, the *motor resonance index* for left-hand conditions (1.33 ± 2.04%) was significantly lower than in baseline (*t*_24_ = –3.21, *p* = .023, *d* = –.64) and from the one obtained after the protocol administration for PAS-conditioned right-hand stimuli (*t*_24_ = –3.02, *p* = .036, *d* = –.6; **Figure 2a**). Here, by looking at raw MEPs, we found that this modulation was due to static trials, which present a significantly higher amplitude than the baseline (**Table 1**).

**Figure 2.**
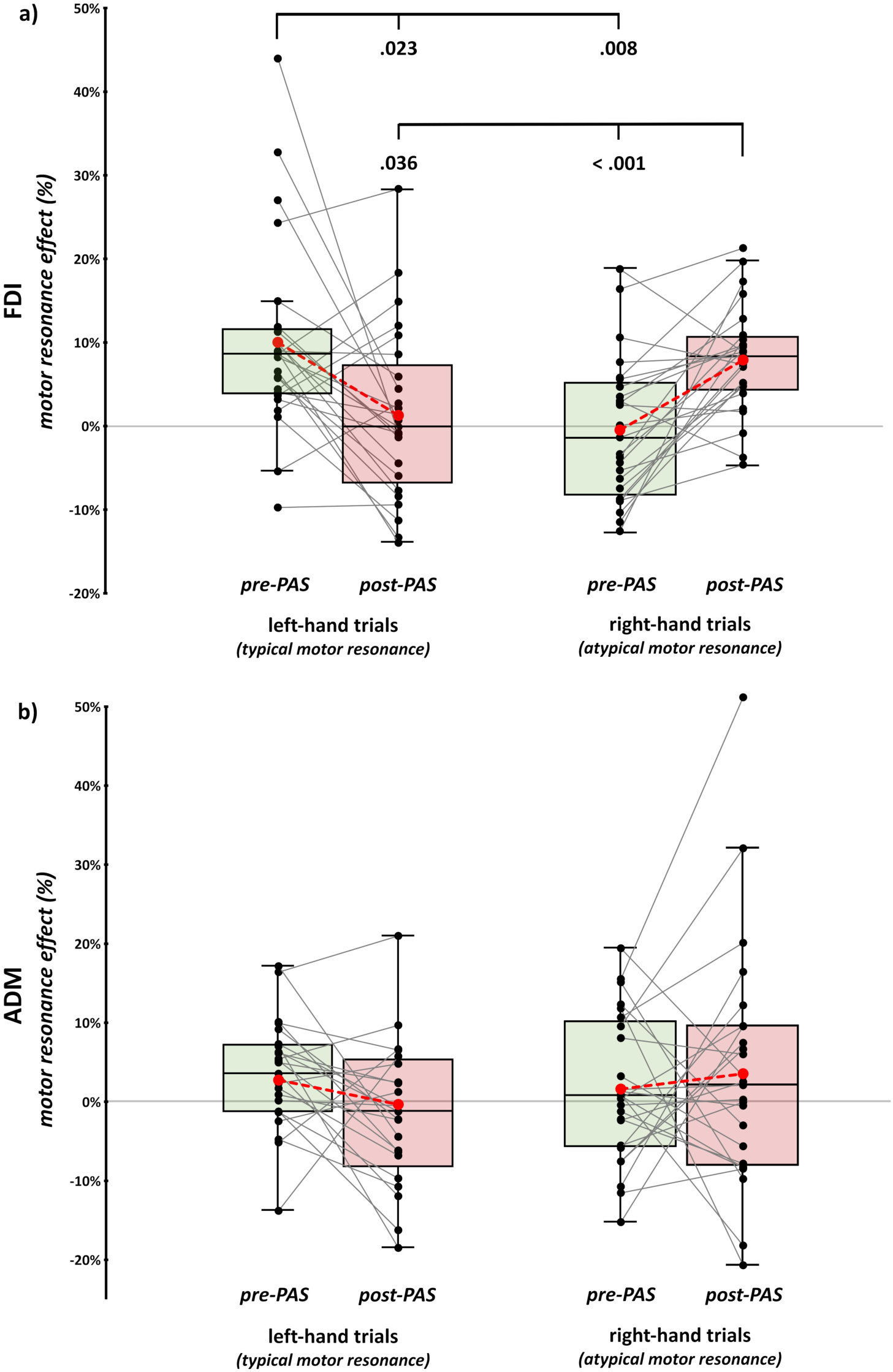
Motor resonance results at the corticospinal level. Motor resonance effects (computed as the ratio between MEP amplitude recorded in movement and static trials) for FDI (**a**) and ADM (**b**) muscles during the observation of action observation task’s left– and right-hand trials before (green boxes) and after (red boxes) the m-PAS administration. In the box-and-whiskers plots, red dots and lines indicate the means of the distributions. The center line denotes their median values. Black dots and grey lines show individual participants’ scores. The box contains the 25^th^ to 75^th^ percentiles of the dataset. Whiskers extend to the largest observation falling within the 1.5 * inter-quartile range from the first/third quartile. Significant *p* values of Bonferroni corrected post-hoc comparisons are reported.

For the ADM muscle – which acts as our control muscle, being not involved in the observed index finger movement – the rmANOVA showed no significant effects of factors ‘viewed Hand’ (*F*_1,24_ < .01, *p* = .965, *η_p_^2^* < .01), ‘Time’ (*F*_1,24_ = .29, *p* = .592, *η_p_^2^* = .01) and their interaction (*F*_1,24_ = .34, *p* = .565, *η_p_^2^* = .01) (**Figure 2b**). This result suggests the muscle-specificity of both typical and PAS-induced motor resonance.

### 3.2. Cortical connectivity signatures of motor resonance during action observation

To investigate PAS-induced changes in interregional connectivity during action observation, we conducted a functional connectivity analysis on TEPs using the wPLI (Vinck, Oostenveld, van Wingerden, Battaglia, & Pennartz, 2011) on source-level activity parcellated into 88 brain regions (for a similar methodology, see: Bianco et al., 2023) comparing connectivity patterns in the different conditions of the action observation task before and after m-PAS administration in the alpha (8-12 Hz) and beta band (13-30 Hz).

#### 3.2.1. Alpha-band connectivity

Overall, TMS-evoked activity while observing unilateral hand movements in our action observation task highlighted widespread M1 alpha connectivity encompassing bilateral frontal, parietal, temporal, and occipital regions (**Figure 3**; see **Supplementary Tables 1, 2** reporting labels of the significant parcels found and **Supplementary Figure 2a** for parcels’ differences between static and movement trials). Before m-PAS administration, observing movements performed by the left hand (i.e., the condition typically eliciting motor resonance) triggered a more distributed network of connections spreading from the right M1, especially towards contralateral frontal regions and the ipsilateral superior temporal gyrus. This cortico-cortical communication may thus represent the mechanism upon which typical motor facilitation relies. In support of this hypothesis, comparing connectivity patterns contingent upon the observation of static and moving hands before and after the m-PAS revealed a marked change. In static trials, m-PAS administration led to greater right M1 connectivity with ipsilateral temporo-occipital parcels, as well as a reduction with bilateral frontal and contralateral parieto-occipital ones. Conversely, in movement trials, there was only a marked decrease in connections and their strength, especially between M1 and frontal parcels of both hemispheres (**Figure 3**). That is to say, the reduction of typical MEP facilitation reliably found after m-PAS administration is mirrored by a whole-brain reorganization of M1 alpha-band connectivity in response to left-hand movement observation.

**Figure 3.**
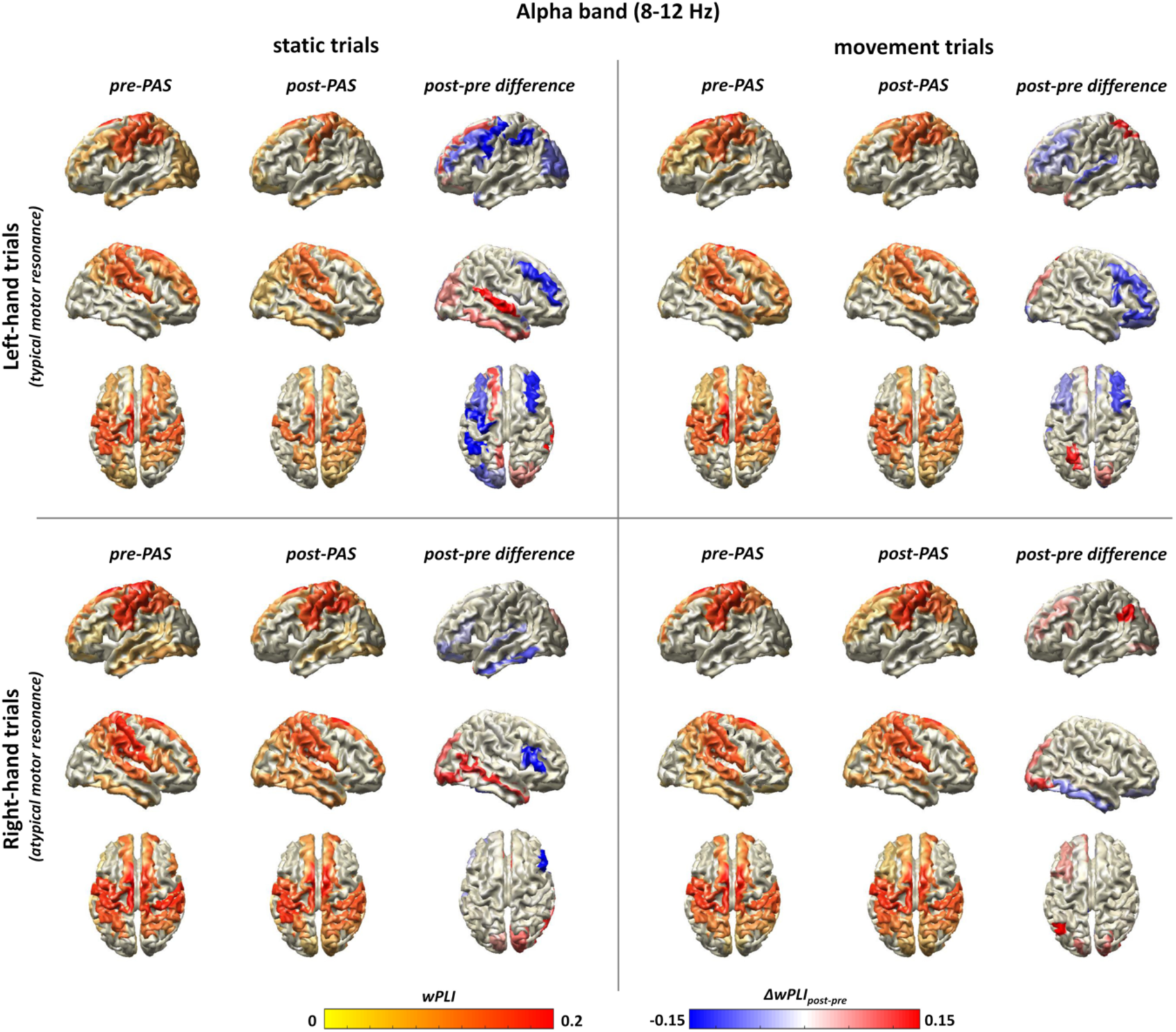
Alpha-band connectivity plots. Plots for alpha-band connectivity (8-12 Hz) after right M1-TMS during static (left panel) and movement (right panel) trials of the action observation task before and after m-PAS administration. Labels of the significant parcels are reported in **Supplementary Tables 1** and **2**.

This evidence is further corroborated by the analysis of the alpha-band *connectivity strength index,* computed for each participant as the sum of all significant connections, in which we reported a significant three-way ‘viewed Hand’ X ‘Visual stimulus’ X ‘Time’ interaction (*F*_1,24_ = 7.34, *p* = .012, *η_p_^2^* = .23). This effect was explored by performing two separate rm-ANOVA for each block of task (i.e., depicting left hand or right hand).

For left-hand conditions, we found a significant effect of factor ‘Time’ (*F*_1,24_ = 24.37, *p* < .001, *η_p_^2^* = .5) and of the interaction ‘Visual stimulus’ X ‘Time’ (*F*_1,24_ = 15.37, *p* <.001, *η_p_^2^* = .39). Post-hoc comparisons revealed that movement trials at baseline showed greater connectivity strength (.8 ± .04) than static ones at both timepoints (*vs.* pre-PAS static trials: .73 ± .04; *t*_24_ = 3.7, *p* = .007, *d* = .74; *vs.* post-PAS static trials: .7 ± .04, *t*_24_ = 4.42, *p* = .001, *d* = .88). Critically, we found a selective reduction of connectivity strength after the m-PAS in trials depicting a movement (.68 ± .04) compared to both baselines (*vs.* pre-PAS static trials: *t*_24_ = – 3.31, *p* = .018, *d* = –.66; *vs.* pre-PAS movement trials: *t*_24_ = –7.01, *p* < .001, *d* = –1.4; **Figure 4a**). Instead, the ‘Visual stimulus’ main effect did not reach statistical significance (*F*_1,24_ = 1.11, *p* = .302, *η_p_^2^* = .04).

**Figure 4.**
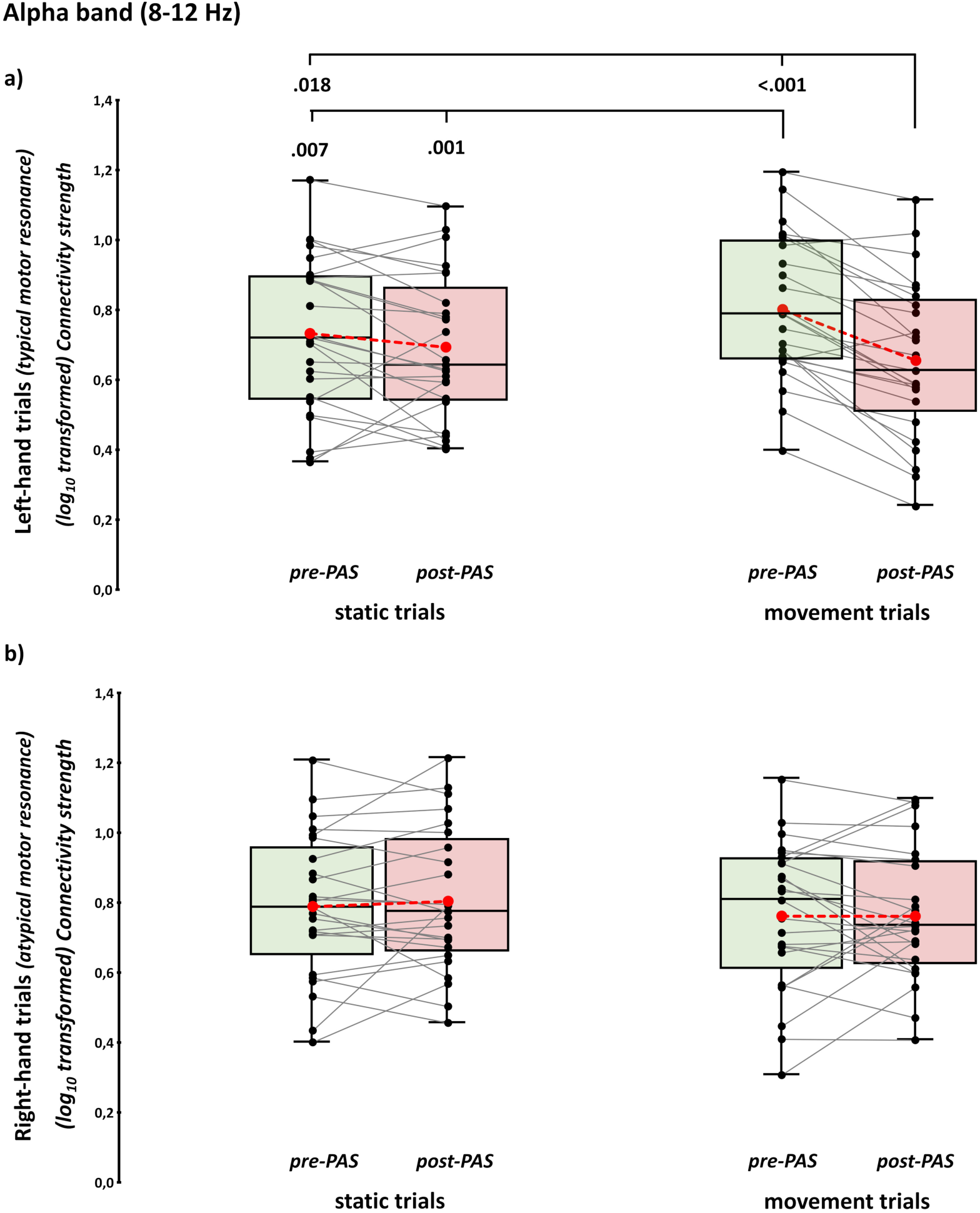
Alpha-band connectivity strength results. (Log_10_ transformed) connectivity strength in the alpha band for left-hand (**a**) and right-hand (**b**) trials of the action observation task before (green boxes) and after (red boxes) m-PAS administration. Raw values were log_10_-transformed to achieve normality of the distribution. In the box-and-whiskers plots, red dots and lines indicate the means of the distributions. The center line denotes their median values. Black dots and grey lines show individual participants’ scores. The box contains the 25^th^ to 75^th^ percentiles of the dataset. Whiskers extend to the largest observation falling within the 1.5 * inter-quartile range from the first/third quartile. Significant *p* values of Bonferroni corrected post-hoc comparisons are reported.

Considering the observation of the right hand, only a significant effect of the main factor ‘Visual stimulus’ was found (*F*_1,24_ = 11.97, *p* = .002, *η_p_^2^* = .33). Regardless of the m-PAS, observing the static hand showed greater connectivity strength values (.8 ± .04) than the movement observation (.76 ± .04). Factor ‘Time’ (*F*_1,24_ = .1, *p* = .752, *η_p_^2^* < .01) and the interaction ‘Visual stimulus’ X ‘Time’ (*F*_1,24_ = .43, *p* = .519, *η_p_^2^* = .02) did not reach statistical significance (**Figure 4b**).

#### 3.2.2. Beta-band connectivity

Beta-band connectivity patterns during action observation (**Figure 5, Supplementary Figure 2b**, and **Supplementary Tables 3, 4**) showed that phase synchronization in this frequency band generally supports M1 connectivity with frontal, parietal, and temporal areas encompassing both the left and right hemispheres. Focusing on connectivity patterns before m-PAS administration, the network activated by the observation of left-hand movements, which gave rise to typical motor resonance, had an overall reduced number of connections, mainly restricted to cortical parcels in the frontal and parietal cortices, compared to the other conditions. Conversely, right-hand conditions showed reduced connectivity with the left precentral and postcentral gyri compared to the left-hand ones. Interestingly, at baseline, both left– and right-hand conditions did not show striking differences in connectivity patterns between movement and static trials (**Supplementary Figure 2b**). Crucially, after m-PAS administration, we reported an opposite pattern of changes with an increase in predominantly posterior connections (i.e., occipital and temporal) during the observation of left-hand movements along with a decrease in frontal and parietal connections during trials depicting right-hand movements, i.e., those conditioned by the m-PAS and associated with the newly acquired motor resonance response. Furthermore, in this latter condition, we also observed increased connectivity with the left pre– and post-central gyri (**Figure 5**). Accordingly, the ‘viewed Hand’ X ‘Visual stimulus’ X ‘Time’ rm-ANOVA on the beta-band *connectivity strength index* indicated a significant three-way interaction (*F*_1,24_ = 52.53, *p* < .001, *η_p_^2^* = .68), suggesting that these patterns are also detectable quantitively.

**Figure 5.**
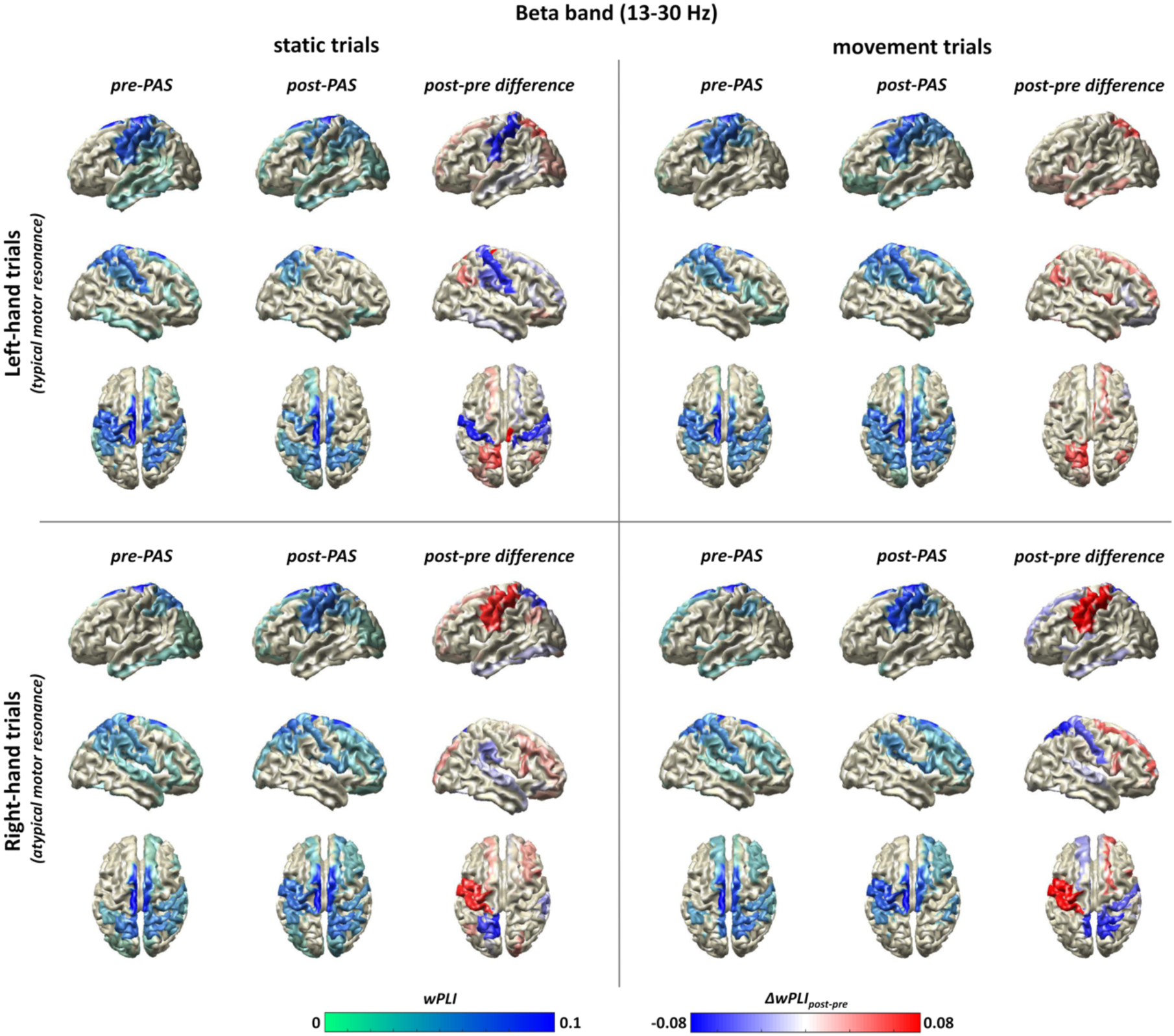
Beta-band connectivity plots. Plots for beta-band connectivity (13-30 Hz) after right M1-TMS during static (left panel) and movement (right panel) trials of the action observation task before and after m-PAS administration. Labels of the significant parcels are reported in **Supplementary Tables 3** and **4**.

Considering the view of the left hand, we found a significant effect of both main factors (‘Visual stimulus’: *F*_1,24_ = 11.15, *p* = .003, *η_p_^2^* = .32; ‘Time’: *F*_1,24_ = 8.88, *p* = .006, *η_p_^2^* = .27) and, crucially, of their interaction (‘Visual stimulus’ X ‘Time’: *F*_1,24_ = 32.81, *p* < .001, *η_p_^2^* = .58). In detail, before m-PAS administration beta-band connectivity strength was lower for movement trials (.35 ± .03) with respect to the observation of static hands (.45 ± .03, *t*_24_ = –6.47, *p* < .001, *d* = –1.3). Conversely, after m-PAS, the observation of movement trials resulted in a greater network strength (.47 ± .04) as compared to static trials (.43 ± .03, *t*_24_ = –3.88, *p* = .004, *d* = –.78; **Figure 6a**). Finally, the connectivity strength of moving trials also differed before vs. after the m-PAS protocol (*t*_24_ = 6.46, *p* < .001, *d* = 1.29).

**Figure 6.**
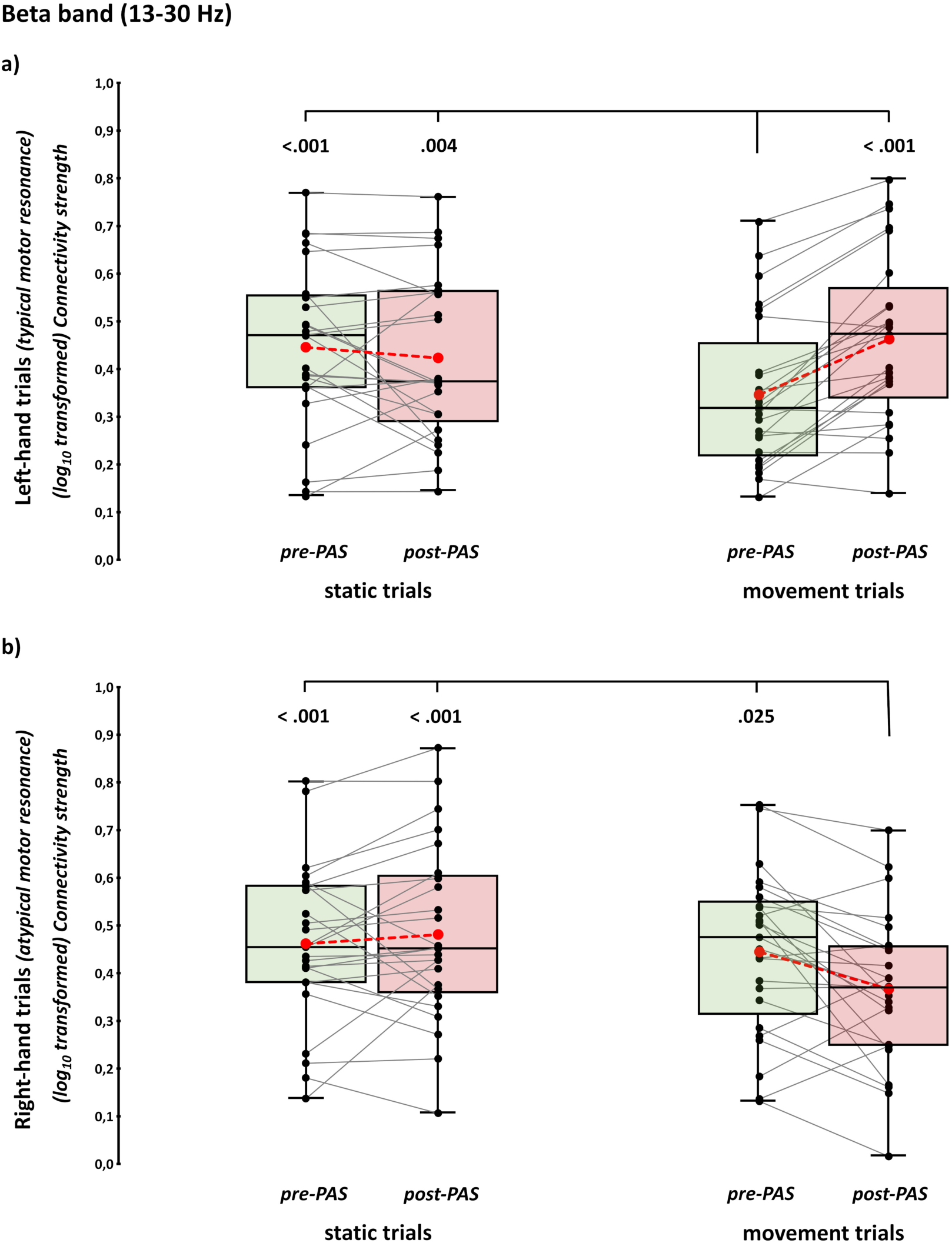
Beta-band connectivity strength results. (Log_10_ transformed) connectivity strength in the beta band for left-hand (**a**) and right-hand (**b**) trials of the action observation task before (green boxes) and after (red boxes) m-PAS administration. Raw values were log_10_-transformed to achieve normality of the distribution. In the box-and-whiskers plots, red dots and lines indicate the means of the distributions. The center line denotes their median values. Black dots and grey lines show individual participants’ scores. The box contains the 25^th^ to 75^th^ percentiles of the dataset. Whiskers extend to the largest observation falling within the 1.5 * inter-quartile range from the first/third quartile. Significant *p* values of Bonferroni corrected post-hoc comparisons are reported.

Concerning the right hand, we found a significant effect of factor ‘Visual stimulus’ (*F*_1,24_ = 26.82, *p* < .001, *η_p_^2^* = .53) and ‘Visual stimulus’ X ‘Time’ interaction (*F*_1,24_ = 20.53, *p* < .001, *η_p_^2^* = .46). Before m-PAS no difference was present between moving (.46 ± .03) and static hands trials (.45 ± .03, *t*_24_ = –.99, *p* > .999, *d* = .2). Conversely, after the administration of the m-PAS observing left-hand movements induced a significant decrease in connectivity strength (.37 ± .03) compared to the observation of static hands (.48 ± .04, *t*_24_ = –6.76, *p* < .001, *d* = –1.35) (**Figure 6b**). Movement trials also differed between before and after the m-PAS protocol (*t*_24_ = –3.17, *p* = .02, *d* = –.63). The factor ‘Time’ did not reach statistical significance (*F*_1,24_ = 26.82, *p* < .001, *η_p_^2^* = .53).

Crucially, no significant differences pre-*vs.* post-protocol administration were found during the observation of both left and right static hands (all *p*s > .05). This pattern of results suggests that PAS-induced changes affected only M1-beta connectivity during action observation, with reverse effects for actions performed by ipsilateral and contralateral hands.

## 4. DISCUSSION

The present study investigated global changes in functional networks underlying typical and newly acquired (i.e., experimentally induced) motor resonance responses to understand how this mechanism is implemented in the brain and the dynamics underpinning its acquisition and modulation. To do so, we computed inter-regional connectivity through TMS-EEG co-registration before and after the administration of the m-PAS, a non-invasive neuromodulatory protocol able to induce novel mirroring phenomenon thanks to Hebbian associative plasticity (Guidali et al., 2020). Besides replicating m-PAS effectiveness in causing atypical motor resonance responses at the corticospinal level, our findings revealed dissociable cortico-cortical signatures of M1-related interregional communication within the alpha and beta bands associated with the occurrence and acquisition of the motor resonance phenomenon.

### 4.1. Emergence of corticospinal motor resonance for PAS-conditioned movement

At the corticospinal level, we replicate the effectiveness of the protocol (Guidali et al., 2020; Guidali, Picardi, Gramegna, et al., 2023), showing that the m-PAS reliably induces Hebbian-like associative plasticity that leads to the transient reshaping of visuomotor representations detectable at the peripheral level, thus resulting in the emergence of motor resonance for ipsilateral (to stimulated hemisphere) hand movements. The effect is muscle-specific (e.g., Avenanti, Bolognini, Maravita, & Aglioti, 2007; Romani, Cesari, Urgesi, Facchini, & Aglioti, 2005; Urgesi, Candidi, Fabbro, Romani, & Aglioti, 2006): the PAS-induced modulations on the motor resonance index is significant only for the FDI muscle, i.e., the effector involved in the movement observed during the m-PAS protocol, but not for the ADM muscle.

Interestingly, we also found a significant decrease in the typical motor resonance response while observing left-hand movements (i.e., contralateral to TMS) after m-PAS administration. This motor resonance ‘shift’ towards the PAS-conditioned movement is similar to modulations reported by previous studies exploiting ‘counter-mirror’ sensorimotor training, which are also accompanied by the loss of MEP facilitation for the muscle originally associated with the observed movement (Catmur, Mars, Rushworth, & Heyes, 2011; Catmur et al., 2007; Cavallo, Heyes, Becchio, Bird, & Catmur, 2014). As noted in the **Results**, differences in raw MEP amplitude data before and after m-PAS administration (see **Table 1**) highlight how the motor resonance reduction for contralateral movements is driven by an increase in MEP amplitude measured during static left-hand conditions, suggesting that this modulation is due to the enhancement of the corticospinal tract output at the sight of static hands, rather than to the inhibition during trials depicting the actual movement. It has to be noted that our previous studies have already reported a trend in this direction, but it never reached statistical significance (see: Guidali et al., 2020). We suggest that the clearcut result of the present work may be related to the higher number of trials planned in our experimental design (80 for each task’s conditions against 20 as in our previous experiments), which was necessary to obtain reliable TMS-EEG measures (Kerwin, Keller, Wu, Narayan, & Etkin, 2018) and may have reduced MEP inter-trials variability (Cuypers, Thijs, & Meesen, 2014), improving the detection of statistically significant effects on this variable.

Overall, MEP results support the evidence that the m-PAS effectively induces sensorimotor learning within the AON by reconfiguring visuomotor representations during action observation of both hand movements. Indeed, the experimentally induced motor resonance response emerged in parallel with a reshaping of the typical phenomenon.

### 4.2. Distinct functional connectivity fingerprints for typical and newly acquired motor resonance

Considering functional connectivity results, we found differential involvement of M1 alpha– and beta-band networks in the occurrence and reshaping of motor resonance for task conditions requiring the observation of the index finger movement. Overall, connectivity analysis applied on TEPs recorded during the action observation task allows us to highlight the involvement of a distributed network connecting M1 to fronto-parietal, temporal, and visual regions oscillating in these two frequency bands (**Figures 3** and **5**). M1 interregional communication in alpha and beta bands could reflect dissociable aspects of human mirroring phenomena, displaying how typical motor resonance is implemented within the motor network and how this mechanism’s plasticity unfolds during the acquisition of a new visuomotor association. This evidence is in line with previous studies highlighting the relevance of alpha and beta rhythms during action observation (Babiloni et al., 2016; Muthukumaraswamy & Johnson, 2004; Qin et al., 2023; Simon & Mukamel, 2016), although most of these investigations primarily focused on regional oscillatory activity instead of addressing oscillatory networks, as in the present work.

#### 4.2.1. Modulation of alpha-band connectivity by m-PAS administration is specific for typical motor resonance

Our results show that M1-related phase synchronization for typical motor resonance is particularly prominent in the alpha band, in line with evidence linking alpha-band phase dynamics to the functional state of M1, both in terms of corticospinal output (Zrenner et al., 2023) and effective connectivity with other cortical regions (Stefanou, Desideri, Belardinelli, Zrenner, & Ziemann, 2018; Zazio, Miniussi, & Bortoletto, 2021). Before m-PAS administration, the M1-related alpha network recruited during the observation of left-hand movements (i.e., the condition triggering typical motor resonance) is overall greater than that recruited during trials depicting the static hand (see **Figure 3**). A more pronounced interregional communication between M1 and the rest of the cortex could benefit corticospinal facilitation, characterizing the typical motor resonance phenomenon as time-established preferential visuomotor representation. Besides being generally linked to the functional status of the motor system, distributed alpha-band functional connectivity has also been related to attentional processes and top-down modulation of sensory activity (Palva & Palva, 2007, 2012). Since attentional and mirroring processes are dissociable but functionally interacting during action observation (Bowman et al., 2017), alpha connectivity could represent a preferential route through which cross-system information grounding the motor resonance phenomenon is integrated, ultimately affecting the corticospinal output. This evidence could explain the more pronounced corticospinal output during the observation of movements performed with the left hand in the baseline. Accordingly, the reshaping of typical motor resonance by m-PAS occurs in parallel with a significant drop in M1 connectivity in the alpha band during the observation of static and moving hands, suggesting that alpha-band dynamics might ground experience-based mirror-like effects in humans.

Nevertheless, if mere interregional alpha-band communication has a deterministic role in the manifestation of motor resonance, we would have expected increased alpha connectivity in the action observation task’s trials depicting the right-hand movement that, after the PAS, elicited the atypical motor facilitation. Instead, we report no significant changes in right-hand conditions after the m-PAS (see **Figure 4b**). Then, it is possible that M1 alpha-band functional networks support motor resonance phenomena established throughout life (Cook et al., 2014), while transient reinforcement of a specific visuomotor representation may not allow for a complete remapping of such mechanism.

#### 4.2.2. Beta-band connectivity patterns of motor resonance reassemble modulations found at the corticospinal level

The pattern of results obtained by analyzing beta-band connectivity strength parallels the one found at the corticospinal level on the *motor resonance index*. Under normal conditions (i.e., before m-PAS), beta-band connectivity during the observation of left-hand movements (i.e., associated with the typical motor resonance phenomenon) is suppressed compared to all other conditions (see **Figure 5**). After m-PAS, the M1-related beta-band network decreased during the observation of PAS-conditioned right-hand movements (i.e., the newly acquired motor resonance response) and increased during the observation of left-hand ones (i.e., no longer associated with corticospinal facilitation after the m-PAS). This evidence suggests that a drop in beta-phase M1 communication may be the critical mechanism supporting the formation of motor resonance by coordinating the activity of motor-related brain regions during action observation.

Beta activity is essential for the functioning of the motor system and control of the corticospinal output (Brittain & Brown, 2014; Engel & Fries, 2010; Wischnewski, Haigh, Shirinpour, Alekseichuk, & Opitz, 2022). Beta oscillations have previously been related to inhibitory neurotransmission in the motor system, especially to gamma-aminobutyric acid (GABA) levels, and proved to be sensitive to plastic changes occurring across the lifespan (Rossiter, Davis, Clark, Boudrias, & Ward, 2014). Furthermore, corticospinal excitability is influenced by beta-phase synchronization, possibly with an opposite relationship compared to ongoing alpha oscillation (Wischnewski et al., 2022). The suppression of beta-band rhythms during action observation and execution is a well-established phenomenon, thus supporting its involvement in these processes (Milston, Vanman, & Cunnington, 2013). Beta-phase synchronization is also supposed to maintain the current motor state and inhibit the initiation of new movements (Engel & Fries, 2010; Picazio et al., 2014). Moreover, beta-band dynamics have been related to the mirroring of touch, which occurred in parallel with a vicarious activation of the somatosensory cortex (Pisoni, Romero Lauro, et al., 2018).

Therefore, a reduction in beta-band connectivity could reflect a release of inhibitory processes similar to what is commonly observed locally at the power level (Barone & Rossiter, 2021). Alternatively, restricting the beta-band oscillatory network around fronto-parietal nodes could signal a functional specialization to promote a more targeted information flow between the motor cortex and premotor and parietal nodes. Indeed, beta-band connectivity is particularly beneficial for communication between premotor and parietal nodes during the action observation as a function of predictability (Qin et al., 2023). In both cases, our data support the idea that M1 interregional dynamics could relate to regulatory mechanisms ruling the activation state of the sensorimotor system during action observation: motor resonance effects at the corticospinal level are sensitive to these dynamics, and their rewriting is reflected in interareal communication networks. While typical mirror-like motor responses exhibit an alpha-band increase and a beta-band decrease in M1 connectivity, newly induced patterns of motor facilitation shaped through visuomotor learning are supported only by the reduction of beta-band M1 cortical networks.

### 4.3. Conclusions and future directions

Our results convincingly present a potential mechanism supporting motor resonance and its plastic modulation after sensorimotor Hebbian-like learning, which appears to be a fundamental process guiding visuomotor matching properties of the human AON (Heyes & Catmur, 2022; Keysers & Gazzola, 2014). Crucially, the experimentally induced visuomotor association partially overlaps M1 interregional communication featured by the typical motor resonance response. Indeed, the newly acquired, m-PAS-induced motor resonance is likely driven by beta-band dynamics resembling those observed for the typical phenomenon in the baseline but lacks the alpha-band modulations that are specific for canonical motor resonance. Newly acquired associations, thus, rely on simpler mechanisms of corticospinal communication, while long-term ones involve additional top-down control encompassing more frontal regions.

The present study also highlights the feasibility of PAS protocols to investigate brain dynamics of high-order cognitive networks, unveiling the cortico-cortical substrates supporting the emergence of motor resonance at the peripheral (corticospinal) level. Such kind of information extends and completes the findings provided by the mere analysis of the corticospinal output, in line with previous TMS-EEG studies showing that M1-TEP and MEP modulations could frame different states of motor cortex activation, thus not implying mutuality in their respective findings (Biabani, Fornito, Coxon, Fulcher, & Rogasch, 2021; Guidali, Zazio, et al., 2023; Madsen et al., 2019). Here, we show that cortical measures like TEPs could be successfully exploited to deepen corticospinal indexes of AON recruitment, providing precious information about the neurophysiological fingerprints of motor resonance and its reshaping following sensorimotor learning.

Furthermore, leveraging the motor resonance phenomenon, our study provides valuable insight for exploiting the m-PAS protocol in clinical settings to induce plasticity within motor regions through AON recruitment. Lastly, recent works have shown how TMS-EEG could be used to refine PAS parameters (i.e., the inter-stimulus interval between paired stimulations) by directly measuring the response delay between TMS perturbation over a cortical target and the following EEG response of the interconnected cortical node (Borgomaneri et al., 2023). In future studies, this approach could also be exploited to optimize PAS-stimulated cortico-cortical pathways, thus better investigating the contribution of different AON cortical nodes (see e.g., Chiappini et al., 2024) to the typical and atypical motor resonance modulations found in the present work.

In conclusion, our results offer the first neurophysiological signature of the complex M1-interareal communication supporting life-established and newly acquired motor resonance responses. The plasticity of the human AON, through PAS-induced sensorimotor associative learning, exploits cortico-cortical connectivity mechanisms that are likely distinct from those established throughout lifetime experience. This is reflected by the evidence that the experimentally induced motor resonance response is not entirely comparable to the typical one, even if the same corticospinal output (i.e., MEP facilitation during action observation) characterizes them.

## ACKNOWLEDGMENTS

We thank Davide Mazzucchelli and Sara Pellegatta for their valuable help in the data collection.

## CRedIT AUTHOR CONTRIBUTION

**Giacomo Guidali:** conceptualization, methodology, investigation, formal analysis, software, data curation, visualization, writing – original draft

**Eleonora Arrigoni:** methodology, investigation, formal analysis, software, visualization, writing – original draft

**Nadia Bolognini:** conceptualization, methodology, supervision, resources, funding acquisition, writing – review & editing

**Alberto Pisoni:** conceptualization, methodology, supervision, validation, resources, writing – review & editing

## SUPPLEMENTARY INFORMATION CAPTIONS

**Supplementary Figure 1**. Plots for the significantly different parcels between movement and static trials in the alpha (8-12 Hz – **a**) and beta-band (13-30 Hz – **b**).

**Supplementary Figure 2**. Sensor level M1-TEP grand average and topography of the main peaks recorded in the eight different experimental conditions (left panels: left-hand blocks; right panels: right-hand blocks; upper row: static trials; lower row: movement trails). C4 electrode, corresponding on average to the stimulation site (i.e., left M1), is depicted in red in the grand average.

**Supplementary Table 1**. AAL labels of the significant parcels (p<.001) found for the alpha band in static and movement trials depicting the left hand, before and after m-PAS administration.

**Supplementary Table 2**. AAL labels of the significant parcels (p<.001) found for the alpha band in static and movement trials depicting the right hand, before and after m-PAS administration.

**Supplementary Table 3**. AAL labels of the significant parcels (p<.001) found for the beta band in static and movement trials depicting the left hand, before and after m-PAS administration.

**Supplementary Table 4**. AAL labels of the significant parcels (p<.001) found for the beta band in static and movement trials depicting the right hand, before and after m-PAS administration.

