## Supplemental materials for "M1 Large-scale Network Dynamics Support Human Motor Resonance and Its Plastic Reshaping"

### - SUPPORTING INFORMATION -

This file contains the following Supplementary Materials:

- Supplementary Figure 1 ………………………………… p. 2

- Supplementary Figure 2 ………………………………… p. 3

- Supplementary Table 1 …………………………………. p. 4

- Supplementary Table 2 …………………………………. p. 7

- Supplementary Table 3 …………………………………. p. 10

- Supplementary Table 4 …………………………………. p. 12

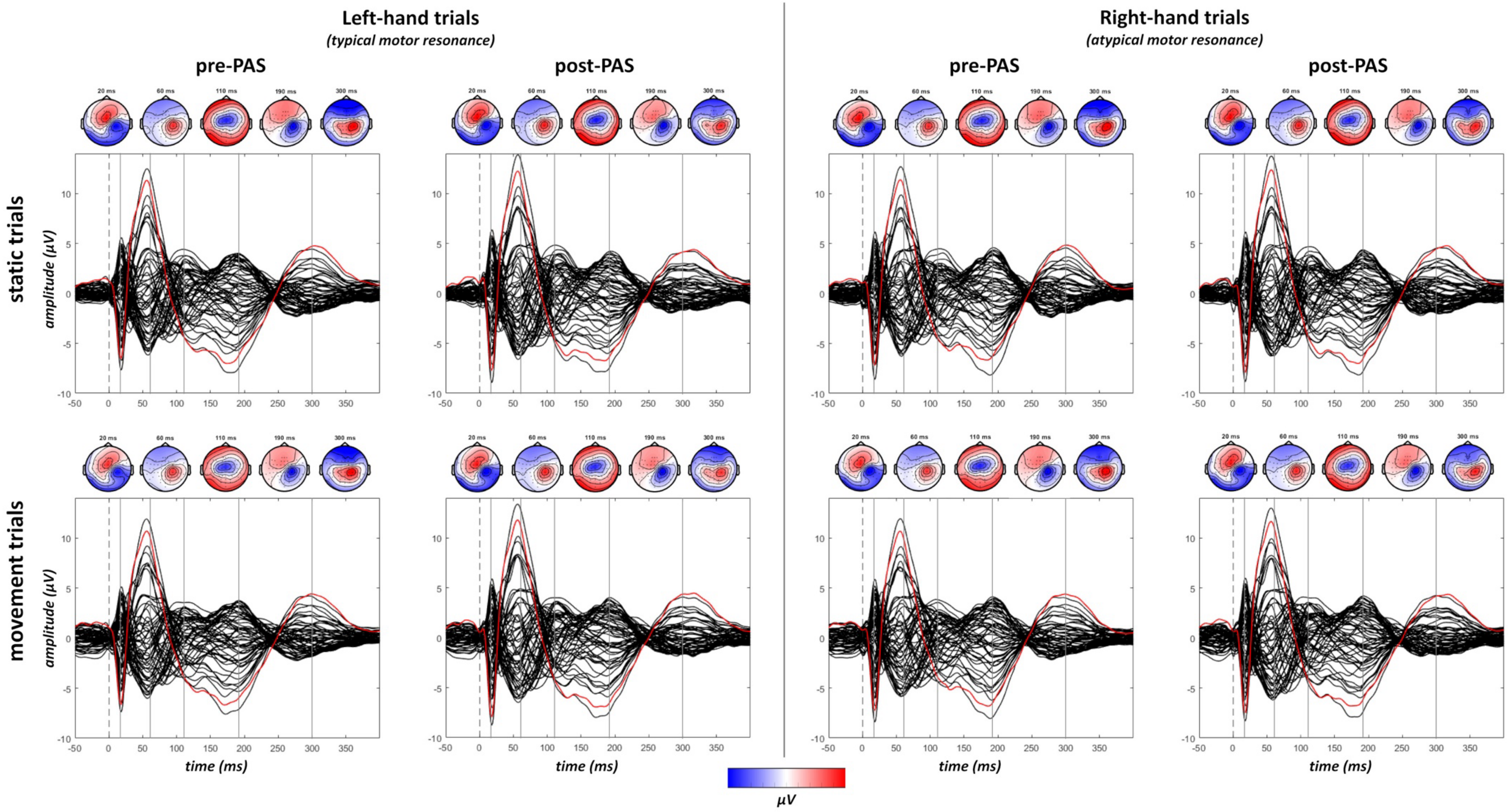

**Supplementary Figure 1**. Sensor level M1-TEP grand average and topography of the main peaks recorded in the eight different experimental conditions (left panels: left-hand blocks; right panels: right-hand blocks; upper row: static trials; lower row: movement trails). C4 electrode, corresponding on average to the stimulation site (i.e., left M1), is depicted in red in the grand average.

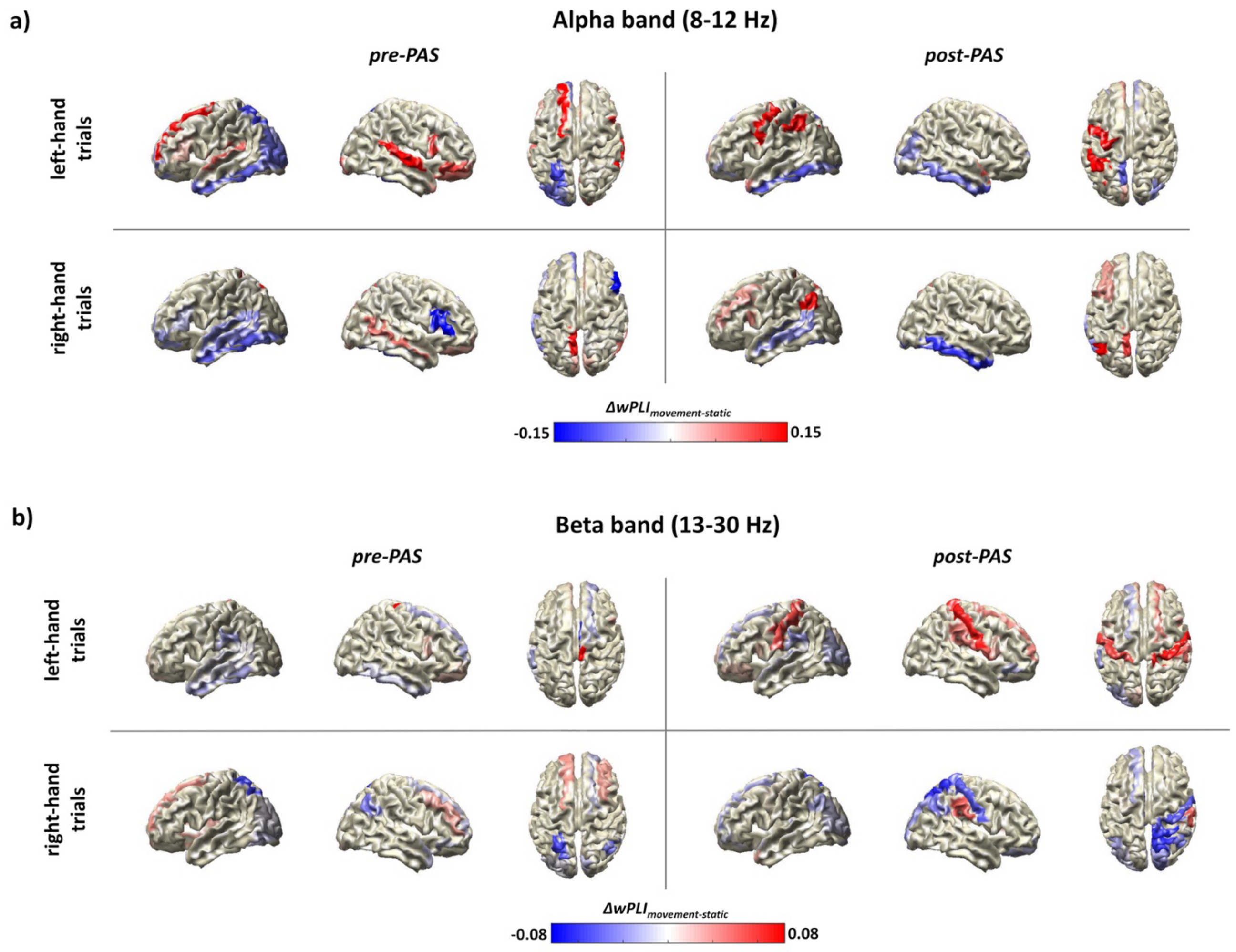

**Supplementary Figure 2.** Plots for the significantly different parcels between movement and static trials in the alpha (8-12 Hz – **a**) and beta-band (13-30 Hz – **b**).

| ALPHA BAND (8-12 Hz) | LEFT-HAND CONDITIONS | | | |
|---|---|---|---|---|
| | **Static trials** | | **Movement trials** | |
| | ***pre-PAS*** | ***post-PAS*** | ***pre-PAS*** | ***post-PAS*** |
| **Frontal lobe** | Precentral Left | Frontal Superior Left | Precentral Left | Precentral Left |
| | Frontal Superior Right | Frontal Superior Right | Frontal Superior Left | Frontal Superior Left |
| | Frontal Superior Orbital Left | Frontal Superior Orbital Left | Frontal Superior Right | Frontal Superior Right |
| | Frontal Middle Left | Frontal Middle Orbital Left | Frontal Superior Orbital Right | Frontal Superior Orbital Left |
| | Frontal Middle Right | Rolandic Operculum Right | Frontal Middle Left | Rolandic Operculum Right |
| | Frontal Inferior Opercular Left | Supplementary Motor Area Left | Frontal Middle Right | Supplementary Motor Area Left |
| | Rolandic Operculum Right | Supplementary Motor Area Right | Frontal Middle Orbital Right | Supplementary Motor Area Right |
| | Supplementary Motor Area Left | Frontal Superior Medial Right | Frontal Inferior Opercular Left | Frontal Superior Medial Left |
| | Supplementary Motor Area Right | Insula Right | Frontal Inferior Opercular Right | Frontal Medial Orbital Left |
| | Frontal Superior Medial Left | Cingulum Middle Left | Frontal Inferior Triangular Left | Insula Right |
| | Frontal Superior Medial Right | Cingulum Middle Right | Frontal Inferior Orbital Right | Cingulum Middle Left |
| | Insula Right | Cingulum Posterior Right | Rolandic Operculum Right | Cingulum Middle Right |
| | Cingulum Middle Left | | Supplementary Motor Area Left | Cingulum Posterior Right |
| | Cingulum Middle Right | | Supplementary Motor Area Right | |
| | | | Olfactory Left | |
| | | | Olfactory Right | |
| | | | Frontal Superior Medial Right | |
| | | | Rectus Right | |
| | | | Insula Right | |
| | | | Cingulum Ant Right | |
| | | | Cingulum Middle Left | |
| | | | Cingulum Middle Right | |
| | | | Cingulum Posterior Right | |
| **Temporal lobe** | Temporal Superior Left | Heschl Left | Heschl Left | Temporal Superior Left |
| | Temporal Pole Superior Right | Temporal Middle Left | Temporal Superior Left | Temporal Pole Superior Right |

| | | | | |
|---|---|---|---|---|
| | Temporal Middle Right | Temporal Middle Right | Temporal Middle Left | Hippocampus Right |
| | Hippocampus Left | Temporal Pole Middle Left | Hippocampus Left | Amygdala Right |
| | ParaHippocampal Left | Hippocampus Left | Hippocampus Right | |
| | Amygdala Left | ParaHippocampal Left | ParaHippocampal Left | |
| | Amygdala Right | Amygdala Right | ParaHippocampal Right | |
| | | | Amygdala Left | |
| | | | Amygdala Right | |
| **Parietal lobe** | Postcentral Left | Postcentral Left | Postcentral Left | Postcentral Left |
| | Postcentral Right | Postcentral Right | Postcentral Right | Postcentral Right |
| | Parietal Superior Left | Parietal Superior Left | Parietal Superior Right | Parietal Superior Left |
| | Parietal Superior Right | Parietal Superior Right | Parietal Inferior Left | Parietal Superior Right |
| | Parietal Inferior Left | Parietal Inferior Right | Parietal Inferior Right | Parietal Inferior Left |
| | Parietal Inferior Right | SupraMarginal Right | SupraMarginal Right | Parietal Inferior Right |
| | SupraMarginal Right | Angular Right | Angular Right | SupraMarginal Right |
| | Angular Right | Precuneus Left | Precuneus Right | Angular Right |
| | Precuneus Right | Precuneus Right | Paracentral Lobule Left | Precuneus Right |
| | Paracentral Lobule Left | Paracentral Lobule Left | Paracentral Lobule Right | Paracentral Lobule Left |
| | Paracentral Lobule Right | Paracentral Lobule Right | | Paracentral Lobule Right |
| **Occipital lobe** | Calcarine Left | Calcarine Right | Calcarine Right | Cuneus Left |
| | Cuneus Left | Cuneus Right | Cuneus Left | Cuneus Right |
| | Lingual Left | Lingual Right | Lingual Left | Occipital Superior Right |
| | Occipital Superior Left | Occipital Superior Right | Lingual Right | |
| | Occipital Middle Left | Occipital Middle Right | Fusiform Left | |
| | Occipital Inferior Left | Occipital Inferior Left | | |
| | Fusiform Left | Fusiform Left | | |
| | | Fusiform Right | | |
| **Subcortical structures** | Caudate Left | Putamen Right | Caudate Left | Putamen Right |
| | Caudate Right | Pallidum Left | Caudate Right | Pallidum Left |
| | Putamen Left | Thalamus Left | Putamen Left | Thalamus Left |

| | | | | |
|---|---|---|---|---|
| | Putamen Right | | Putamen Right | Heschl Left |
| | Pallidum Left | | Pallidum Left | |
| | Thalamus Left | | Thalamus Left | |
| | | | Thalamus Right | |

**Supplementary Table 1.** AAL labels of the significant parcels (p<.001) found for the alpha band in static and movement trials depicting the left hand, before and after m-PAS administration.

| ALPHA BAND (8-12 Hz) | RIGHT-HAND CONDITIONS | | | |
|---|---|---|---|---|
| | **Static trials** | | **Movement trials** | |
| | ***pre-PAS*** | ***post-PAS*** | ***pre-PAS*** | ***post-PAS*** |
| **Frontal lobe** | Precentral Left | Precentral Left | Precentral Left | Precentral Left |
| | Frontal Superior Left | Frontal Superior Left | Frontal Superior Left | Frontal Superior Left |
| | Frontal Superior Right | Frontal Superior Right | Frontal Superior Right | Frontal Superior Right |
| | Frontal Superior Orbital Left | Rolandic Operculum Right | Frontal Superior Orbital Right | Frontal Middle Left |
| | Frontal Middle Orbital Left | Supplementary Motor Area Left | Rolandic Operculum Right | Frontal Inferior Opercular Left |
| | Frontal Inferior Opercular Right | Supplementary Motor Area Right | Supplementary Motor Area Left | Rolandic Operculum Right |
| | Frontal Inferior Triangular Left | Frontal Superior Medial Left | Supplementary Motor Area Right | Supplementary Motor Area Left |
| | Frontal Inferior Triangular Right | Frontal Superior Medial Right | Olfactory Right | Supplementary Motor Area Right |
| | Rolandic Operculum Right | Insula Right | Frontal Superior Medial Right | Frontal Superior Medial Left |
| | Supplementary Motor Area Left | Cingulum Anterior Left | Rectus Right | Frontal Superior Medial Right |
| | Supplementary Motor Area Right | Cingulum Anterior Right | Insula Right | Insula Right |
| | Frontal Superior Medial Left | Cingulum Middle Left | Cingulum Anterior Right | Cingulum Anterior Left |
| | Frontal Superior Medial Right | Cingulum Middle Right | Cingulum Middle Left | Cingulum Anterior Right |
| | Insula Right | Cingulum Posterior Right | Cingulum Middle Right | Cingulum Middle Left |
| | Cingulum Middle Left | | Cingulum Posterior Left | Cingulum Middle Right |
| | Cingulum Middle Right | | Cingulum Posterior Right | Cingulum Posterior Right |
| | Cingulum Posterior Right | | | |
| **Temporal lobe** | Heschl Left | Heschl Left | Heschl Left | Heschl Left |
| | Temporal Superior Right | Temporal Superior Right | Temporal Pole Superior Left | Temporal Pole Superior Left |
| | Temporal Middle Right | Temporal Pole Superior Left | Temporal Pole Middle Left | |
| | Temporal Pole Middle Left | Temporal Middle Left | Hippocampus Right | |
| | Hippocampus Left | Temporal Pole Middle Left | ParaHippocampal Right | |
| | Hippocampus Right | Hippocampus Right | Amygdala Right | |
| | ParaHippocampal Left | ParaHippocampal Left | | |
| | ParaHippocampal Right | ParaHippocampal Right | | |
| | Amygdala Right | Amygdala Right | | |

| | | | | |
|---|---|---|---|---|
| **Parietal lobe** | Postcentral Left | Postcentral Left | Postcentral Left | Postcentral Left |
| | Postcentral Right | Postcentral Right | Postcentral Right | Postcentral Right |
| | Parietal Superior Left | Parietal Superior Left | Parietal Superior Left | Parietal Superior Left |
| | Parietal Superior Right | Parietal Superior Right | Parietal Superior Right | Parietal Superior Right |
| | Parietal Inferior Left | Parietal Inferior Left | Parietal Inferior Left | Parietal Inferior Left |
| | Parietal Inferior Right | Parietal Inferior Right | Parietal Inferior Right | Parietal Inferior Right |
| | SupraMarginal Right | SupraMarginal Right | SupraMarginal Right | SupraMarginal Right |
| | Angular Right | Angular Right | Angular Right | Angular Left |
| | Precuneus Right | Precuneus Right | Precuneus Left | Angular Right |
| | Paracentral Lobule Left | Paracentral Lobule Left | Precuneus Right | Precuneus Left |
| | Paracentral Lobule Right | Paracentral Lobule Right | Paracentral Lobule Left | Precuneus Right |
| | | | Paracentral Lobule Right | Paracentral Lobule Left |
| | | | | Paracentral Lobule Right |
| **Occipital lobe** | Calcarine Left | Calcarine Left | Calcarine Left | Calcarine Left |
| | Lingual Left | Calcarine Right | Cuneus Left | Calcarine Right |
| | Occipital Inferior Left | Cuneus Left | Cuneus Right | Cuneus Left |
| | Fusiform Left | Cuneus Right | Lingual Left | Cuneus Right |
| | Fusiform Right | Lingual Left | Lingual Right | Lingual Left |
| | | Lingual Right | Fusiform Right | Occipital Superior Left |
| | | Occipital Superior Left | | Occipital Superior Right |
| | | Occipital Superior Right | | Occipital Inferior Left |
| | | Occipital Inferior Left | | Occipital Inferior Right |
| | | Occipital Inferior Right | | |
| | | Fusiform Left | | |
| | | Fusiform Right | | |
| **Subcortical structures** | Putamen Right | Putamen Right | Caudate Left | Putamen Right |
| | Pallidum Left | Pallidum Left | Caudate Right | Pallidum Left |
| | Thalamus Left | Thalamus Left | Putamen Left | Thalamus Left |
| | Thalamus Right | | Putamen Right | |
| | | | Pallidum Left | |

| | | | |
|---|---|---|---|
| | | | Thalamus Left |

**Supplementary Table 2.** AAL labels of the significant parcels ($p<.001$) found for the alpha band in static and movement trials depicting the right hand, before and after m-PAS administration.

| BETA BAND (13-30 Hz) | LEFT HAND CONDITIONS | | | |
|---|---|---|---|---|
| | **Static trials** | | **Movement trials** | |
| | ***pre-PAS*** | ***post-PAS*** | ***pre-PAS*** | ***post-PAS*** |
| **Frontal lobe** | Precentral Left | Precentral Left | Precentral Left | Precentral Left |
| | Frontal Superior Right | Frontal Superior Left | Frontal Superior Orbital Left | Frontal Superior Right |
| | Frontal Inferior Triangularis Right | Frontal Superior Orbital Left | Frontal Superior Orbital Right | Frontal Superior Orbital Left |
| | Supplementary Motor Area Left | Frontal Inferior Orbital Right | Frontal Middle Orbital Right | Frontal Middle Orbital Left |
| | Supplementary Motor Area Right | Supplementary Motor Area Left | Frontal Inferior Opercular Right | Frontal Inferior Opercular Left |
| | Olfactory Left | Supplementary Motor Area Right | Frontal Inferior Triangularis Right | Frontal Inferior Opercular Right |
| | Olfactory Right | Olfactory Left | Supplementary Motor Area Left | Frontal Inferior Orbital Left |
| | Frontal Medial Orbital Right | Insula Left | Supplementary Motor Area Right | Rolandic Operculum Right |
| | Insula Right | Insula Right | Olfactory Left | Supplementary Motor Area Left |
| | Cingulum Anterior Right | Cingulum Middle Left | Frontal Superior Medial Left | Supplementary Motor Area Right |
| | Cingulum Middle Left | Cingulum Middle Right | Frontal Medial Orbital Right | Frontal Superior Medial Left |
| | Cingulum Middle Right | Cingulum Posterior Left | Rectus Left | Insula Left |
| | Cingulum Posterior Right | Cingulum Posterior Right | Rectus Right | Insula Right |
| | | | Insula Right | Cingulum Middle Left |
| | | | Cingulum Anterior Left | Cingulum Middle Right |
| | | | Cingulum Anterior Right | Cingulum Posterior Left |
| | | | Cingulum Middle Left | Cingulum Posterior Right |
| | | | Cingulum Posterior Right | |
| **Temporal lobe** | Temporal Superior Right | Heschl Right | | Temporal Pole Superior Right |
| | Temporal Pole Superior Right | Temporal Pole Superior Right | | Temporal Middle Left |
| | Temporal Middle Left | Temporal Middle Left | | Temporal Middle Right |
| | Temporal Middle Right | Temporal Middle Right | | Hippocampus Right |
| | Temporal Pole Middle Left | Hippocampus Left | | ParaHippocampal Right |
| | Hippocampus Right | Amygdala Left | | Amygdala Right |
| | ParaHippocampal Left | Amygdala Right | | |
| | ParaHippocampal Right | | | |
| | Amygdala Left | | | |

| | | | | |
|---|---|---|---|---|
| | Amygdala Right | | | |
| **Parietal lobe** | Postcentral Left | Parietal Superior Left | Postcentral Left | Postcentral Left |
| | Postcentral Right | Parietal Superior Right | Postcentral Right | Postcentral Right |
| | Parietal Superior Right | Parietal Inferior Left | Parietal Superior Right | Parietal Superior Left |
| | Parietal Inferior Left | Parietal Inferior Right | Parietal Inferior Left | Parietal Superior Right |
| | Parietal Inferior Right | SupraMarginal Left | Parietal Inferior Right | Parietal Inferior Left |
| | SupraMarginal Left | Angular Right | SupraMarginal Right | Parietal Inferior Right |
| | SupraMarginal Right | Precuneus Left | Precuneus Right | SupraMarginal Right |
| | Precuneus Right | Precuneus Right | Paracentral Lobule Left | Angular Right |
| | Paracentral Lobule Left | Paracentral Lobule Left | Paracentral Lobule Right | Precuneus Left |
| | | Paracentral Lobule Right | | Precuneus Right |
| | | | | Paracentral Lobule Left |
| | | | | Paracentral Lobule Right |
| | | | | Paracentral Lobule Right |
| **Occipital lobe** | Lingual Left | Calcarine Left | | Calcarine Left |
| | Fusiform Right | Occipital Middle Left | | Cuneus Left |
| | | Fusiform Right | | Occipital Superior Left |
| | | | | Fusiform Right |
| **Subcortical structures** | Caudate Left | Caudate Right | Putamen Left | Putamen Left |
| | Putamen Left | Putamen Right | | Pallidum Left |
| | Putamen Right | Pallidum Right | | |
| | Pallidum Left | Thalamus Right | | |
| | Pallidum Right | | | |
| | Thalamus Right | | | |

**Supplementary Table 3.** AAL labels of the significant parcels ($p<.001$) found for the beta band in static and movement trials depicting the left hand, before and after m-PAS administration.

| BETA BAND (13-30 Hz) | RIGHT-HAND CONDITIONS | | | |
|---|---|---|---|---|
| | **Static trials** | | **Movement trials** | |
| | ***pre-PAS*** | ***post-PAS*** | ***pre-PAS*** | ***post-PAS*** |
| **Frontal lobe** | Frontal Superior Right | Precentral Left | Frontal Superior Left | Precentral Left |
| | Frontal Inferior Triangularis Right | Frontal Superior Left | Frontal Middle Right | Frontal Superior Right |
| | Frontal Inferior Orbital Right | Frontal Superior Right | Frontal Inferior Triangularis Right | Frontal Middle Right |
| | Rolandic Operculum Right | Frontal Superior Orbital Right | Rolandic Operculum Right | Frontal Middle Orbital Right |
| | Supplementary Motor Area Left | Frontal Middle Right | Supplementary Motor Area Left | Frontal Inferior Opercular Right |
| | Supplementary Motor Area Right | Frontal Middle Orbital Left | Supplementary Motor Area Right | Frontal Inferior Triangularis Right |
| | Olfactory Left | Frontal Middle Orbital Right | Olfactory Left | Rolandic Operculum Right |
| | Olfactory Right | Frontal Inferior Opercular Right | Olfactory Right | Supplementary Motor Area Left |
| | Frontal Superior Medial Right | Frontal Inferior Triangularis Right | Frontal Superior Medial Left | Supplementary Motor Area Right |
| | Rectus Right | Rolandic Operculum Right | Frontal Superior Medial Right | Olfactory Right |
| | Insula Right | Supplementary Motor Area Left | Frontal Medial Orbital Left | Insula Right |
| | Cingulum Middle Left | Supplementary Motor Area Right | Rectus Left | Cingulum Middle Right |
| | Cingulum Middle Right | Rectus Left | Insula Left | |
| | Cingulum Posterior Left | Rectus Right | Insula Right | |
| | Cingulum Posterior Right | Insula Right | Cingulum Anterior Left | |
| | | Cingulum Anterior Left | Cingulum Middle Left | |
| | | Cingulum Middle Left | Cingulum Middle Right | |
| | | Cingulum Middle Right | Cingulum Posterior Left | |
| | | Cingulum Posterior Left | Cingulum Posterior Right | |
| | | Cingulum Posterior Right | | |
| **Temporal lobe** | Heschl Left | Heschl Right | Heschl Left | Temporal Pole Superior Right |
| | Temporal Superior Left | Hippocampus Left | Temporal Pole Superior Right | Hippocampus Left |
| | Temporal Pole Superior Right | Hippocampus Right | Temporal Middle Right | Hippocampus Right |
| | Temporal Middle Left | ParaHippocampal Left | Hippocampus Right | ParaHippocampal Right |
| | Temporal Middle Right | ParaHippocampal Right | Amygdala Left | Amygdala Left |
| | Hippocampus Right | | | |

| | | | | |
|---|---|---|---|---|
| | ParaHippocampal Left | | | |
| | ParaHippocampal Right | | | |
| | Amygdala Left | | | |
| **Parietal lobe** | Postcentral Right | Postcentral Left | Postcentral Right | Postcentral Left |
| | Parietal Superior Left | Postcentral Right | Parietal Superior Right | Parietal Inferior Left |
| | Parietal Superior Right | Parietal Superior Right | Parietal Inferior Left | Parietal Inferior Right |
| | Parietal Inferior Left | Parietal Inferior Left | Parietal Inferior Right | SupraMarginal Right |
| | Parietal Inferior Right | Parietal Inferior Right | SupraMarginal Right | Paracentral Lobule Left |
| | SupraMarginal Right | Angular Left | Precuneus Left | Paracentral Lobule Right |
| | Angular Right | Angular Right | Precuneus Right | |
| | Precuneus Left | Precuneus Right | Paracentral Lobule Left | |
| | Precuneus Right | Paracentral Lobule Left | Paracentral Lobule Right | |
| | Paracentral Lobule Left | Paracentral Lobule Right | | |
| | Paracentral Lobule Right | | | |
| **Occipital lobe** | Calcarine Left | Calcarine Left | Lingual Right | |
| | Cuneus Right | Calcarine Right | Fusiform Right | |
| | Occipital Superior Left | Cuneus Right | | |
| | Occipital Middle Left | Lingual Left | | |
| | Occipital Inferior Left | Occipital Superior Right | | |
| | Fusiform Left | Occipital Middle Left | | |
| | Fusiform Right | | | |
| **Subcortical structures** | Caudate Right | Caudate Right | Caudate Left | Caudate Right |
| | Putamen Left | Putamen Right | Caudate Right | Putamen Left |
| | Thalamus Left | Pallidum Left | Putamen Left | Putamen Right |
| | | Pallidum Right | Thalamus Left | Pallidum Left |

**Supplementary Table 4.** AAL labels of the significant parcels (p<.001) found for the beta band in static and movement trials depicting the right hand, before and after m-PAS administration.
